# Adaptation to low parasite abundance affects immune investment strategy and immunopathological responses of cavefish

**DOI:** 10.1101/647255

**Authors:** Robert Peuß, Andrew C. Box, Shiyuan Chen, Yongfu Wang, Dai Tsuchiya, Jenna L. Persons, Alexander Kenzior, Ernesto Maldonado, Jaya Krishnan, Jörn P. Scharsack, Brian P. Slaughter, Nicolas Rohner

## Abstract

Reduced parasite infection rates in the developed world are suspected to underlie the rising prevalence of autoimmune disorders. However, the long-term evolutionary consequences of decreased parasite exposure on an immune system are not well understood. We used the Mexican tetra *Astyanax mexicanus* to understand how loss of parasite diversity influences the evolutionary trajectory of the vertebrate immune system by comparing river with cave morphotypes. Here, we present field data that affirms a strong reduction in parasite diversity in the cave ecosystem and show that cavefish immune cells display a more sensitive proinflammatory response towards bacterial endotoxins. Surprisingly, other innate cellular immune responses, such as phagocytosis, are drastically decreased in cavefish. Using two independent single-cell approaches, we identified a shift in the overall immune cell composition in cavefish as the underlying cellular mechanism, indicating strong differences in the immune investment strategy. While surface fish invest evenly into the innate and adaptive immune system, cavefish shifted immune investment to the adaptive immune system, and here, mainly towards specific T-cell populations that promote homeostasis. Additionally, inflammatory responses and immunopathological phenotypes in visceral adipose tissue are drastically reduced in cavefish. Our data indicate that long term adaptation to low parasite diversity coincides with a more sensitive immune system in cavefish, which is accompanied by a reduction of the immune cells that play a role in mediating the proinflammatory response.

## Main text

Important efforts in hygiene and medical treatment in most industrialized countries have reduced microbial and parasitic infections considerably^1^. While this indisputably improves health and increases life expectancy, the diverse effects on the immune system are not well understood. Host-parasite interactions are the major driving force in the evolution of the immune system^2^. From an evolutionary perspective, the maintenance and control of the immune system is associated with costs^2,3^, so a reduction in parasite diversity should theoretically increase the fitness of the host.

Interestingly, the opposite has been observed. Based on a number of studies, it has been hypothesized that decreased exposure to parasites or biodiversity in general has contributed to the rising numbers of autoimmune diseases in the developed world^4-7^. This phenomenon has been described as the “Old Friends hypothesis”^8^, which argues that the reactivity of the vertebrate immune system depends on the exposure to macroparasites (e.g., helminths) and microparasites (e.g., bacteria, fungi and viruses). These parasites, that the host has coevolved with, are necessary for the host to develop a proper functional immune system and to minimize autoimmune reactions that potentially result in immunopathology (e.g. type 1 diabetes or artheriosclerosis)^8^.

Despite important insights into the physiological underpinnings of autoimmune diseases ^9^, we still lack fundamental knowledge of how autoimmune diseases initially develop. Human populations have only been confronted with this decreased parasite diversity for a couple of generations - very recently in evolutionary terms. This raises the question of how the immune system adapts to such environmental changes in the long term. Given the significant impact on fitness of autoimmune disorders^9^, evolutionary adaptations of the immune system to environments with low biodiversity and thereby low parasite diversity^10,11^ are likely to have been deployed.

The vertebrate immune system is composed of two main systems, the innate and the adaptive immune system. The innate immune system is essential for the initial response against pathogens. Given the short lifetime and high complexity, innate immune cells, such as granulocytes, are thought to be very costly for the host^12^. The adaptive immune system of vertebrates is defined by its long-term protection against pathogens (e.g. through the production of pathogen-specific antibodies). Cells of the adaptive immune systems, such as B- and T-cells, are thought to be less costly for the host due to their low complexity and longevity^12^.

Given the differences in costs, it has been suggested that the vertebrate immune system is capable of adjusting its immune investment strategy^13-15^. The host can invest to different degrees into the innate or adaptive immune cells depending on the parasite abundance in the host environment^13-15^. Accordingly, these different immune investment strategies result in specifc differences in the immune responses^16^.

Based on these phenotypic responses, adaption to environments with low parasite abundance should result in a fixed immune investment strategy that is optimized for host fitness. To explore this idea, we utilized an eco-immunological approach in the Mexican tetra *Astyanax mexicanus*, to study how local adaptation of one host species to environments with a stark difference in parasite diversity affects the immune system of the host.

There are cave and surface adapted populations of this species that have adapted to their respective environments for approximately 50-200 thousand years^17,18^. One important hallmark of cave environments is an overall decrease in biodiversity, including parasite diversity^19,20^. Here we present field data from a cavefish population (Pachón) and one surface fish population (Río Choy), which confirms a stark difference in macroparasite abundance between these two habitats and indicates a higher immune activity of surface fish compared to cavefish under natural conditions. Both, cavefish and surface fish populations can be bred and raised for generations in the lab under identical environmental conditions, which readily facilitates the identification of heritable changes. Therefore, we used lab populations that were derived from the wildtype Pachón and Río Choy populations and an additional cavefish population (Tinaja) to investigate immunological consequences deriving from adaptational processes to environments with low parasite diversity. We demonstrate that cavefish immune cells display a more sensitive and prolonged immune response of proinflammatory cytokines towards bacterial endotoxins *in vitro,* similar to other vertebrate host species in environments with low biodiversity^21,22^. Using an image-based immune cell clustering approach (Image3C) and single cell RNA sequencing (scRNAseq) we show that the observed differences in the cellular immune responses are accompanied by differences in the immune investment strategy, where cavefish produce more lymphoid cells (adaptive immunity) than myeloid cells (innate immunity). We demonstrate that the altered immune investment strategy does not generally affect all lymphocytes but mainly leads to an overrepresentation of T-cells in cavefish.

Notably, we found that a large proportion of overrepresented T-cells in cavefish is represented by *γδ*T-cells. This T-cell population is known for its regulatory role in several autoimmune diseases^23^ and the ability to recognize foreign antigens in a MHC (major histocompatibility complex)-independent manner, thereby bridging the gap between the innate and adaptive immune system^24^. Further scRNAseq analysis of the acute inflammatory response in fish treated with lipopolysaccharides (LPS) revealed transcriptional changes in innate and adaptive immune cells as well as in hematopoietic stem cells (HSCs) that may drive the observed changes in the immune investment strategy. In addition, we observed differences in the adaptive response of T- and B-cells, where cavefish display a higher activation response than surface fish. Finally, we show that the reduction of granulocytic and monocytic cells in cavefish leads to reduced immunopathological consequences for visceral fat storage, which has been described as an adaptional response towards low food supply in the cave environment^25,26^.

We started our investigation by collecting wild fish in their natural habitat to study the differences in parasite abundance between river and cave environments. The differences in parasite abundance between river and cave habitats of *A. mexicanus* have not been studied in detail and are mainly based on assumptions that derive from theoretical models^10^. We collected 16 surface fish (Río Choy) and 16 cavefish (Pachón), respectively (Fig. 1a), and examined them for parasite infections as described before^27^. We found varying numbers of endo- and ectoparasites in surface fish (Fig. 1b, see also Figure S1). Interestingly, we did not detect macroparasite infections in the sampled cavefish (Fig. 1b). While we cannot exclude the possibility of viral or bacterial infections in the cavefish population, we did not detect any obvious signs of systemic or tissue specific infections during the parasitological examination. The lower parasite infection rate in wild cavefish is also reflected in a significant lower spleen somatic index (an elevated immune activity in fish coincides with a swelling of the spleen and increases spleen somatic index^28^) in wild cavefish compared to the surface fish samples (Fig. 1c; mean spleen somatic index in surface fish of 0.663 vs. mean spleen somatic index in cavefish of 0.304; *p* = 0.0047, One-way ANOVA). Given the strong impact of host – parasite interaction on the evolution of the immune system^2^, we speculated that these extreme differences in parasite abundance between cavefish and surface fish environment result in functional and/ or physiological changes to the cavefish immune system.

**Figure 1:**
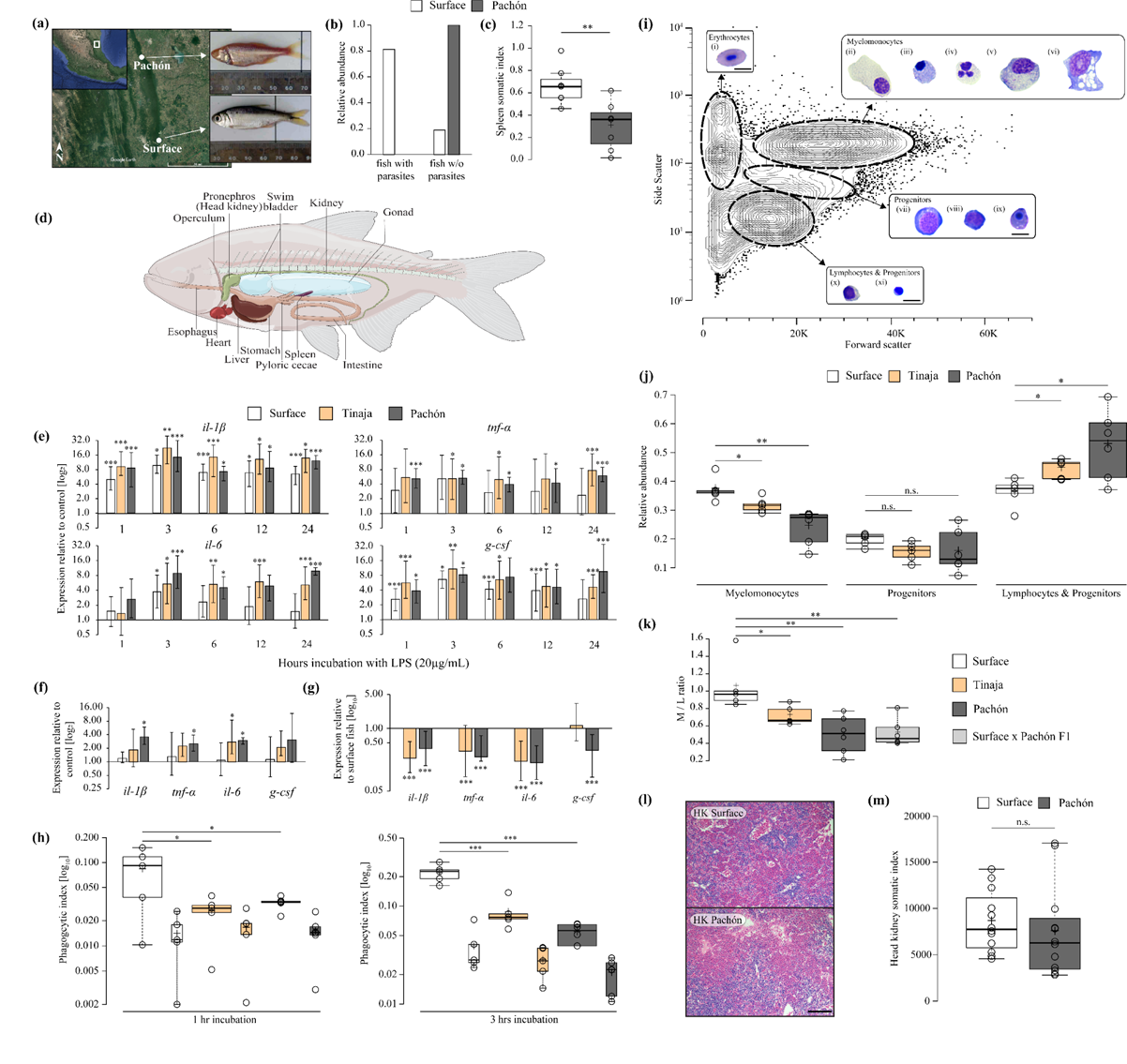
Adaptation to river and cave habitats with stark differences in parasite diversity result in changes of the proinflammatory response and immune investment strategy. (a) Collection sites of *A. mexicanus* surface fish (Río Choy) and cavefish (Pachón). (**b**) Number of fish with and without visible ecto- and endoparasites. (**c**) Immune activity in wild surface and cavefish (n=7 for each) using the spleen somatic index, that is calculated by (weight [mg] (spleen) / weight [mg] (fish)) x 1,000. Significances were determined using a one-way ANOVA. (d) Cartoon of adult *A*. *mexicanus* indicates anatomical position of the main hematopoietic and lymphoid organ, the head kidney (HK) that was used for subsequent in vitro experiments from surface fish and cavefish lab strains. (**e-f**) RT-qPCR analysis of pro-inflammatory cytokines, interleukin-1beta (*il-1β*), tumor necrosis factor alpha (*tnf-α*), interleukin-6 (*il-6*) and granulocyte colony stimulating factor (*g-csf*), of HK cells from surface fish and cavefish after incubation with (e) 20 µg/mL lipopolysaccharide (LPS) at various timepoints or (f) 0.2 µg/mL LPS after 24 hours relative to HK cells incubated without LPS for the given time point. Plotted is the mean of three independent experiments with standard error (SE) on a log2 scale. For all RT-qPCR results, PBS control samples from each time point and sample were used as the reference to calculate relative expression of target genes for each timepoint and fish, respectively. (**g**) RT-qPCR based expression analysis of proinflammatory cytokines *il-1β*, *tnf-α*, *il-6*, *g-csf* of cavefish relative to surface fish of naïve HK samples across all timepoints as shown in (e) (n=18). Significance values were determined by a pairwise fixed reallocation randomization test using REST2009 software. (**h**) Box plot presentation of relative phagocytic rate of HK cells from surface fish and cavefish incubated with Alexa-488 coupled *Staphylococcus aureus*. To control for passive uptake of Alexa-488 coupled *S. aureus*, a control sample was incubated with Alexa-488 coupled *S. aureus* in the presence of 80µg cytochalasin B (CCB) and its phagocytic rate is presented in small boxes of the same color of the respective fish population and timepoint. Significant differences between surface fish (n=5), Tinaja (n=5) and Pachón (n=6) for each timepoint were determined by two-way Anova (see Table S2 for statistical details). (**i**) Representative contour plot of HK cells from three surface *A. mexicanus* after FACS analysis using scatter characteristics showing 99 % of all events. Four different populations (erythrocytes, myelomonocytes, progenitors and lymphocytes & progenitors) were identified and sorted for May-Grünwald Giemsa staining. Images of representative cells that were found in each population are shown for each population and identified based on similar approaches in zebrafish^31,32^ as (i) mature erythrocytes, (ii) promyelocytes, (iii) eosinophiles, (iv) neutrophiles, (v) monocytes, (vi) macrophages, (vii) erythroblasts, (viii) myeloblasts, (ix) erythroid progenitors, (x) lymphocytes and (xi) undifferentiated progenitors. Scale bar is 10 µm. (**j**) Box plots of relative abundances of HK cells from surface and cavefish within the immune populations as defined by scatter characteristics in (**i**). Significances between surface (n=5), Tinaja (n=5) and Pachón (n=6) were determined by one-way ANOVA and subsequent FDR. (**k**) Box plot representation of myelomonocyte / lymphocyte (M / L) ratio from surface (n=5), Tinaja (n=5), Pachón (n=6) and surface x Pachón F1 hybrids (n=6). Significances were determined by one-way ANOVA and subsequent FDR. (**l**) H & E stained section of the head kidney from surface fish and Pachón cavefish (scale bar is 100 µm). (**m**) HK somatic index (mean number of head kidney cells per body weight [mg] fish) is shown for n=12 fish for surface fish and Pachón cavefish. Testing for significant differences was done using one-way ANOVA. For all box plots; center lines show the medians, crosses show means; box limits indicate the 25th and 75th percentiles as determined by R software; whiskers extend 1.5 times the interquartile range from the 25th and 75th percentiles, data points are represented by circles. Significances are indicated as * for p ≤ 0.05; ** for p ≤ 0.01 and *** for p ≤ 0.001 for all experiments.

To explore immunological differences in the potential to develop immunopathological phenotypes between surface fish and cavefish, we first investigated the proinflammatory immune response, which generally precedes immunopathological phenotypes^29^. To trigger such a proinflammatory response we used bacterial endotoxins, LPS, in cultures with extracted leukocytes. We focused on the the pronephros (head kidney, HK) (Figure 1d), the main hematopoietic and lymphoid organ, and a site of antigen representation in teleost fish ^30^. We incubated head kidney cells from surface, Pachón and Tinaja fish with LPS (20 µg/mL) for 1, 3, 6, 12 and 24 hours, respectively and measured gene expression of the proinflammatory cytokines *il-1β*, *tnf-α*, *il-6 and g-csf* in relation to control samples (saline (PBS)) using RT-qPCR (Figure 1e, see Table S1 for details). Head kidney cells from cavefish populations showed an overall greater inducible response upon LPS treatment than head kidney cells from surface fish *in vitro* over time (Fig. 1e). Specifically, only the gene expression of *il-1β* remained significantly elevated in surface fish after 24 hours (Figure 1e). In contrast, cavefish expression of all tested proinflammatory cytokines remained significantly upregulated after 24 hrs. Since the cavefish response was saturated at this LPS concentration, we repeated the analysis with a 100x fold lower LPS exposure (Fig. 1f). Here, LPS treated head kidney cells from surface fish no longer displayed a significant response of any of the proinflammatory cytokines, while Pachón cavefish cells still showed significant expression for *il-1β*, *tnf-α and il-6* compared to untreated cells (Fig. 1f). This increased sensitivity was not present to the same degree in the Tinaja cave population, since we only found an increase in the expression of *il-6* (Fig. 1f).

This increased sensitivity of cavefish head kidney cells towards LPS *in vitro* is supported by previous findings of an increased immune and scarring response after wounding of the Pachón cavefish compared to surface fish^33^. However, the observed differences in the proinflammatory response could be strongly affected by differences in the number of cells that produce these proinflammatory cytokines. To account for this, we directly compared baseline expression of *il-1β*, *tnf-α*, *il-6 and g-csf* in naïve head kidney cells of Pachón and Tinaja to surface fish (Fig. 1g). Surprisingly, the expression of all tested proinflammatory cytokines was significantly reduced in Pachón cavefish samples relative to surface fish cells (Fig. 1g). In the case of the proinflammatory cytokine *il-1β*, for example, naïve Pachón cavefish head kidney cells produced 61 % less transcript than surface fish cells (relative expression Pachón vs. surface fish *il-1β* = 0.383, *p* ≤ 0.001*, pairwise fixed reallocation randomization test*, Fig. 1g). Similar to the Pachón cavefish population, the Tinaja cave population differed in the expression of *il-1β*, *tnf-α* and *il-6* compared to surface fish but not in the expression of *g-csf* (Fig. 1g). Here it is noteworthy that while we only obtained parasite data from one cavefish population, we reasoned that different cave habitats share similar environmental features. To test whether functional differences of the immune system also appear in other cavefish populations we included, where it was feasible, a second, independently derived, cavefish population (Tinaja) in the experimental setup.

Given the difference in cytokine expression upon LPS exposure, we wanted to test whether other cellular immune functions, such as phagocytosis, differ between cavefish and surface fish. We conducted a phagocytosis experiment, in which we quantified the ability of head kidney cells to phagocytize Alexa-488 tagged *Staphylococcus aureus* cells (Thermo Fisher) *in vitro* at different timepoints (Fig. 1h, see Fig. S2 for gating strategy). Using pairwise comparison, we found a significant decrease of the phagocytic rate of both cavefish populations at both timepoints compared to surface fish (Fig. 1h, see Data File 1 for detailed statistic report).

The decreased baseline expression of the proinflammatory cytokines and the decreased phagocytosis rate in cavefish could be the result of changes in the immune cell composition, since both of these cellular immune functions are mainly fulfilled by cells with a myelomonocytic origin (such as granulocytes and monocytes) in teleost fish^32,34,35^. To assess immune cell composition, we analyzed scatter information from head kidney derived single cell suspensions from surface, Tinaja and Pachón fish. Using similar analytical approaches previously described for zebrafish^31,32^, we identified four distinct cell cluster: an erythroid, a myelomonocyte, a progenitor and a lymphoid/progenitor cluster (Fig. 1i). To confirm the identity of these clusters, head kidney cells from each cluster were sorted, and cells were stained with May-Grünwald Giemsa stain. Based on comparative morphological analysis^31,32^, we identified (i) erythrocytes, (ii) promyelocytes, (iii) eosinophiles, (iv) neutrophiles, (v) monocytes, (vi) macrophages, (vii) erythroblasts, (viii) myeloblasts, (ix) erythroid progenitors, (x) lymphocytes and (xi) undifferentiated progenitors (i.e. hematopoietic stem cells, common lymphoid progenitors and common myeloid progenitors) within the four cell clusters (Fig. 1i). When we compared relative abundances of the three immune cell clusters, we identified fewer myelomonocytic cells in both cavefish populations (mean relative abundance of cells in myelomonocyte cluster in surface fish is 0.37 vs. 0.32 in Tinaja, *p* ≤ 0.05, and vs. 0.24 in Pachón, *p* ≤ 0.01; one-way ANOVA, FDR corrected; Fig. 1j) and an increased number of cells in the lymphocyte & progenitor cluster (mean relative abundance of cells in lymphocyte & progenitor cluster in surface fish is 0.36 vs. 0.44 in Tinaja, *p* ≤ 0.05, and vs. 0.53 in Pachón, *p* ≤ 0.05; one-way ANOVA; FDR corrected, Fig. 1j). We used these relative abundances to calculate the myelomonocyte / lymphocyte (M / L) ratio. This ratio is an indication of an individuals relative investment in either innate (myelomonocyte) or adaptive (lymphocyte) immune cell populations^12,36^.

We found profound differences in the immune investment strategy between surface fish and cavefish. While surface fish have a relatively balanced investment in myelomonocyte and lymphoid immune cells, cavefish invest less into myelomonocytic cells than into lymphoid immune cell populations (mean M / L ratio in surface fish is 1.06 vs. 0.72 in Tinaja, *p* ≤ 0.05, and vs. 0.50 in Pachón, *p* ≤ 0.01; one-way ANOVA, FDR corrected, Fig. 1k). Since all fish were raised under identical laboratory conditions, the observed differences in the immune investment strategy point towards a genetic basis for this trait. To further study this, we analyzed surface x Pachón hybrids and found a similar M / L ratio as in the parental Pachón population, indicating that the change in the immune investment strategy of the Pachón population is a dominant trait (mean M/L ratio in surface x Pachón F1 of 0.52, Fig. 1k).

Given the strong difference in parasite abundance and resource availibillity^25^ between cave and surface environments, cavefish potentially benefited from a change in the immune investment that reduces the resource allocation into the immune system (see McDade, et al. ^12^ for review on costs of innate and adaptive immune defences). To rule out the possibility that changes in the head kidney morphology and / or total numbers of cells in the head kidney are responsible (and could potentially compensate) for the observed differences in the M/L ratio, we compared morphology and total cell numbers of the head kidney from surface fish and Pachón cavefish. We observed no general differences in cell morphology (see Fig. 1l) and no significant changes in the absolute cell number from the entire head kidney between the fish populations (mean absolute cell number of entire head kidney per mg fish weight for surface fish was 8556 and 7460 for cavefish, *p* = 0.53, one-way ANOVA; Fig. 1m).

To identify more specific differences in the immune cell composition of cavefish and surface fish, we clustered head kidney cells based on cell morphological features using Image3C^37^. This tool uses image-based flow cytometry and advanced clustering algorithms to cluster cells based on their morphology and cellular features such as granularity of the cytoplasm or nucleus, independent of an observer bias that has been reported for such analysis^38^. This makes it an effective method for organisms lacking established transgenic lines or antibodies to identify specific immune cell populations.

First, we sorted myelomonocyte, lymphocyte and progenitor cell populations as identified in Fig. 1i in order to reduce mature erythrocytes from the single cell suspensions of head kidney cells (see Methods for details). In total, we recorded 10,000 cells by image cytometry from each replicate surface fish (Río Choy, n=5) and cavefish (Pachón, n=6) (Fig. 2a; see methods for more details) and identified 21 distinct cell clusters (Fig. 2b). The identity of each cluster was determined based on cell image galleries from each cluster (see Data File 2 for complete cell gallery) in comparison to the histological staining of sorted cells as presented in Fig. 1i. In addition, to verify certain cellular features (e.g., complexity of nuclei, cell shape, see Table S4 for feature details) within a certain cluster, we used a feature intensity/cluster correlation analysis (Fig. S3). Clusters were assigned to one of the following categories based on their morphological features: myelomonocytes (relatively large cells with medium to high granularity, irregular shaped nuclei and high cytoplasm to nuclei ratio; cluster 2, 11, 13, 14, 16); lymphocytes/progenitors (relatively small cells with low granularity and low cytoplasm to nuclei ratio; cluster 4, 7, 9, 18, 19) and mature erythrocytes/doublets/debris (cluster 1, 3, 5, 6, 8, 10, 12, 15, 17, 20, 21) (Fig. 2b). In line with the scatter analysis, we found a significant reduction of cells within the myelomonocyte category in cavefish compared to surface fish (mean relative abundance of 0.468 cells in surface fish vs 0.320 cells in cavefish; *p* ≤ 0.001, one-way Anova and FDR correction, Fig. 2c).

**Figure 2:**
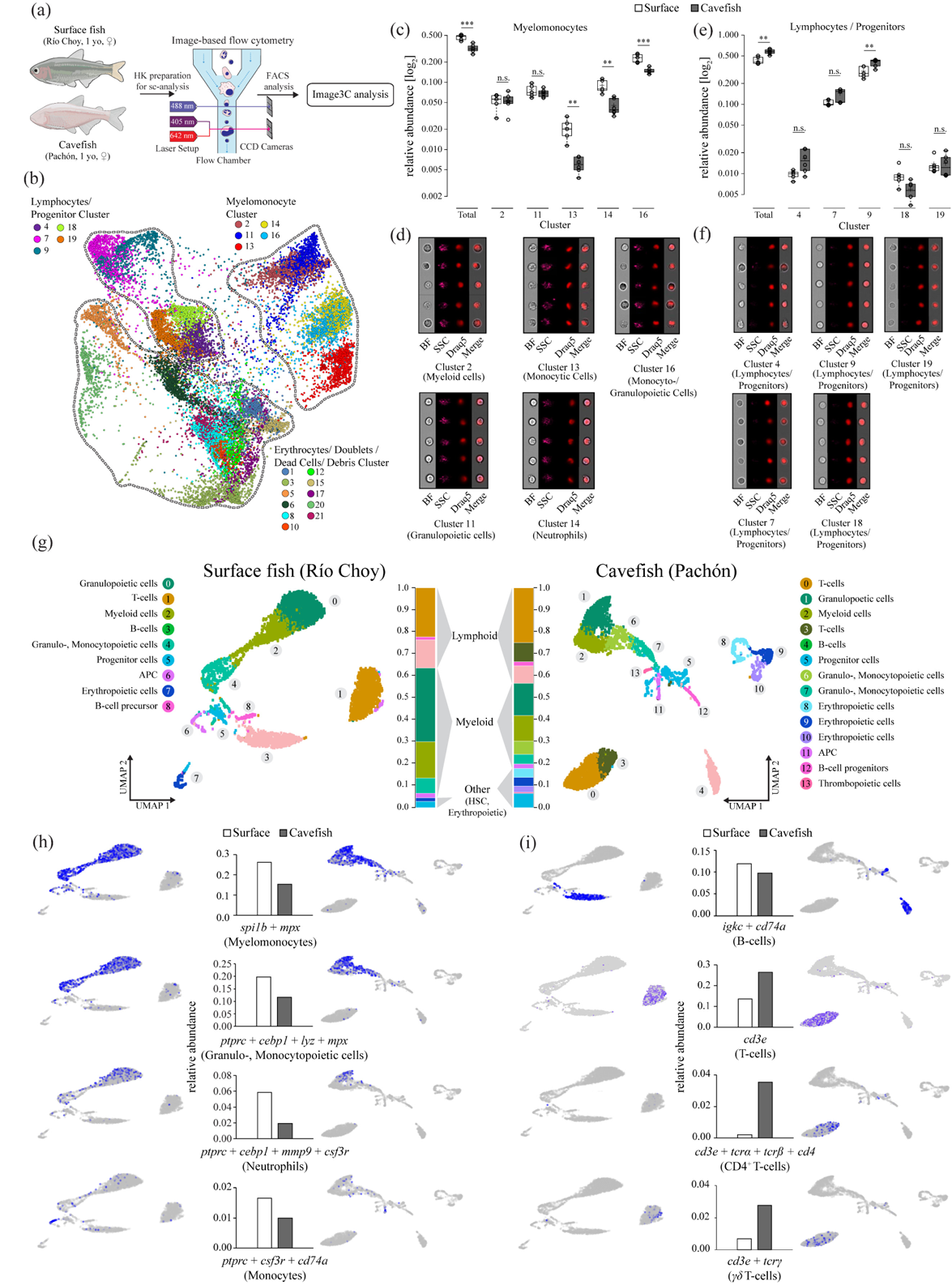
Cell composition analysis of the head kidney of *A. mexicanus* surface and cave morphotypes using cell morphological and genetic features. (**a**) Experimental setup for semi-unsupervised cell morphological clustering using the Image3C pipeline. (**b**) Force directed layout graph of cell clusters based on morphological feature intensities from HK cells of surface fish (n=5) and Pachón cavefish (n=6). Each dot represents a cell and each color represents a unique cluster. Clusters were combined into three categories (Myelomonocytes, Lymphocytes/Progenitors, mature Erythrocytes/Doublets/Debris) based on their morphology (see Data File 1 for cell galleries for each cluster). (**c**) Relative abundances of cells within each cluster of the myelomonocyte category with (**d**) image galleries of each cluster. (**e**) Relative abundances of cells within each cluster of lymphocyte/progenitor with (**f**) image galleries of each cluster. Significant differences in the relative abundance of cells within each category (total) or cluster were determined by one-way ANOVA and subsequent FDR. Significance values are indicated as ** for p ≤ 0.01 and *** for p ≤ 0.001. For box plots; center lines show the medians, crosses show means; box limits indicate the 25th and 75th percentiles as determined by R software; whiskers extend 1.5 times the interquartile range from the 25th and 75th percentiles, data points are represented by circles. Cell galleries show images of brightfield (BF), side scatter (SSC), nuclei (visualized through nuclei dye Draq5) and merged image of BF and Draq5 (Merge). (**g**) UMAP plot of single cell RNA sequencing analysis from one surface fish (Río Choy) and cavefish (Pachón) head kidney, respectively, where each dot represents a cell and each color represents unique cell cluster as shown in the legend. Overall relative abundances are given for each cluster shown in the respective color. (**h-i**) Relative abundance of specific cell populations of surface fish and cavefish and their location within UMAP representation of specific cell types from (h) myelomonocytes and (i) lymphocytes based on the expression of given gene(s). See Data File 3 for gene enrichment in each cluster.

More specifically, we identified differences in the relative abundance of monocytic cells (Fig. 2c-d; relatively large cells with complex structured cytoplasm, high granularity and kidney shaped nuclei; cluster 13; mean relative abundance of 0.021 cells in surface fish vs 0.006 cells in Pachón cavefish; *p* ≤ 0.01; one way ANOVA, FDR corrected), neutrophils (Fig. 2c-d; medium sized cells with evenly distributed cytoplasm, high granularity and multi-lobed nuclei; cluster 14; mean relative abundance of 0.088 cells in surface fish vs 0.044 cells in Pachón cavefish; *p* ≤ 0.01 one way ANOVA, FDR corrected) and monocytic-, granulocytic- and promyelocytic cells (Fig. 2c-d; medium to large cells with high granularity; cluster 16; mean relative abundance of 0.231 cells in surface fish vs 0.148 cells in Pachón cavefish; *p* ≤ 0.01, one way ANOVA, FDR corrected) between surface fish and cavefish.

The reduction of almost all myeloid cell populations suggests that there is an overall reduced investment in the innate immune system in Pachón cavefish. The resulting reduction of granulocytes and monocytes in cavefish head kidney is in line with the observed decreased baseline expression of proinflammatory cytokines and phagocytic rate. Furthermore, we found that cells within the lymphocyte/progenitor category (Fig. 2e-f) are generally overrepresented in cavefish when compared to surface fish (mean relative abundance of cells within lymphocytes/progenitor category: surface fish 0.433 vs. cavefish 0.580; *p* ≤ 0.01 one-way Anova, FDR corrected, Fig. 2e).

Most cluster in the lymphocytes/progenitor category did not differ significantly between surface fish and cavefish with the exception of cluster 9 (surface fish 0.295 vs. cavefish 0.399; *p* ≤ 0.01 one-way Anova, FDR corrected, Fig. 2e), which is the most abundant cluster in this category. Here it is noteworthy, that the M/L ratio we obtained for surface and cavefish with the Image3C approach is similar to the M/L ratio we obtained before (Fig. 1k) using standard scatter information (M/L ratio surface 1.10 vs. 0.56 in Pachón, *p* ≤ 0.001 one-way Anova, Fig. S4). However, given the morphological similarities (see Data File 2) of early progenitor cells of hematopoietic lineages and specific lymphocyte cell types (B-cells, T-cells), we were not able to further resolve the identity of these cluster. Therefore, we took a genetic approach. We performed single-cell RNA sequencing of head kidney cells, where mature erythrocytes were removed from the head kidney cell suspension through FACS sorting as described above. We used one female adult surface fish (Río Choy) and one age, size and sex-matched cavefish (Pachón). Using 10x genomics, we captured 5874 surface fish HK cells and 4717 cavefish HK cells representing most hematopoietic cell linages (Fig. 2g). Cluster identification was done using a comparative approach using gene expression data from other teleost fish species (for details see Methods, Data File 3 for overall gene enrichment in each cluster). Consistent with the morphological analyses, we found an overall reduction in all cells of the myeloid linage in cavefish compared to surface fish. In detail, cluster analysis revealed the reduction of myeloid cells (relative abundance of *spi1b* (*pu.1) + mpx* cells in surface fish 0.221 vs. 0.132 in cavefish, Fig. 2h) and granulo- and monocytopoietic cells (relative abundance of (*ptprc + cebp1 + lyz + mpx* cells in surface fish 0.167 vs. 0.100 in Pachón cavefish, Fig. 2h). Furthermore, we verified the reduction of mature neutrophils (relative abundance of *ptprc* (*cd45*) + *cebp1* + *mmp9* cells in surface fish 0.0586 vs. 0.0191 in Pachón cavefish, Fig. 2h) and monocytes (relative abundance of *ptprc* (*cd45*) + *csf3r* + *cd74a* cells in surface fish 0.0165 vs. 0.010 in Pachón cavefish, Fig. 2h). Importantly, the analysis of the lymphoid cell linage revealed that there are distinct differences in specific lymphoid cell populations between cavefish and surface fish and not an overall increase in lymphoid cells in cavefish as the morphological analysis might suggest (Fig. 2i).

While we found an over-representation in the relative abundance of lymphocytes in cavefish compared to surface fish (Fig. 2g), we found almost identical relative abundances of B-lymphocytes (relative abundance of *igkc + cd74a* cells in surface fish 0.119 vs. 0.098 in Pachón cavefish). In contrast, we found clear differences in the numbers of HK resident T-cells (relative abundance of *cd3e* cells in surface fish 0.136 vs. 0.265 in Pachón cavefish, Fig. 2i). We identified increased numbers of CD4+ T-cells in cavefish (relative abundances of *cd3e* + tcr*α* + tcr*β + cd4-1* in surface fish 0.002 vs 0.031 in Pachón cavefish, Fig. 2i). Interestingly, we observed that *γ^+^δ^+^*CD4^−^ CD8^-^ (*γδ*) T-cells reside in higher proportions in the headkidney of cavefish than in surface fish (relative abundances of *cd3e* + *tcrγ* in surface fish 0.007 vs. 0.028 in Pachón cavefish, Fig. 2i). *γδ* T-cells are a lymphoid cell population that potentially functions as a bridge between the innate and adaptive immune system due to its ability to recognize antigens in an MHC-independent manner^24^ and has only been discovered recently in other teleost species^39^. Furthermore, *γδ* T-cells are reported to play a significant role in the development of autoimmune diseases^23^ and in homeostasis and inflammation of mammalian adipose tissue^40^.

The changes in the immune investment strategy of cavefish suggests that the inflammatory response of Pachón cavefish could be affected, since numbers of cells that drive proinflammatory responses (monocytes and neutrophils) are decreased and cells that can promote homeostatsis (*γδ* T-cells) are increased in Pachón cavefish. In addition, the dominance of the Pachón cavefish immune investment phenotype suggests that there are genetic differences that drive these changes. To address these questions we designed another scRNA-seq experiment (Fig. 3a). We injected surface fish (Río Choy) and cavefish (Pachón) with either PBS or LPS and dissected the head kidney 3 hours after injection (Fig. 3a). We removed the majority of mature erythrocytes through FACS sorting as described before. Considering an unique molecular identifier count of ≥ 500, we obtained a mean of 12,128 cells for each of the eight samples with a mean number of 667 genes per cell in each sample (Fig. 3a). With the increased numbers of cells per sample we were able to resolve cellular identities at higher resolution as shown in Fig. 2g. As done before, we mainly used cell-specific expression data from zebrafish to identify the identity of the single cluster (for details see Methods section). We find higher numbers of myeloid cells in both treatment groups of surface fish, which also resulted in higher numbers of granulopoietic (granulocytes and their precursor cells) and monocytopoietic (monocytes and their precursor cells) cell cluster (Fig. 3b). We were not able to identify a specific cluster of eosinophils, but this is mainly due to the lack of a suitable genetic marker for this cell type. In contrast to the increased numbers of myeloid cells in surface fish, we found an increased number of T-cells in both treatment groups of cavefish similar to the previous experiment (see Fig. 3b and Fig. 2g). Again, the increased numbers of cells per sample enabled a better resolution of the different T-cell populations (Fig. 3b). Based on the gene expression profile, we found naïve T-cells, CD8 + T-cells, Treg cells, CD4+ T-cells, and *γδ* T-cells in cavefish, while we only found naïve and CD4+ T-cells in surface fish (Fig. 3b and 3c). Even though we find T-cells with the same identity in surface fish, the low numbers probably prevented clustering of these cells into a unique T-cell cluster in surface fish. This underlines the differences in the immune investment strategy between surface fish and Pachón cavefish, where surface fish invest more into innate immune cells and Pachón cavefish more into adaptive, more specifically T-cells, immune cells.

**Figure 3:**
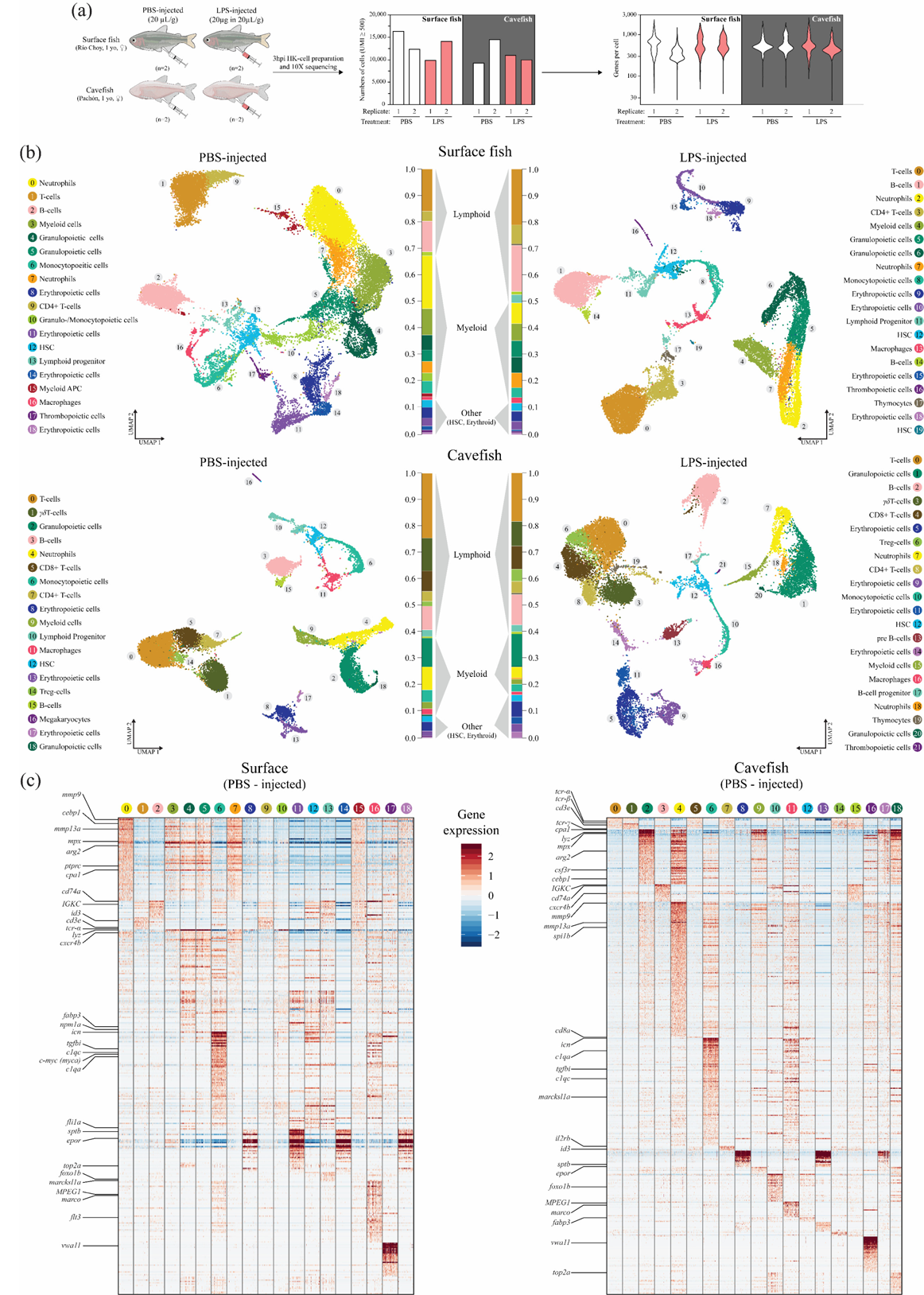
Cellular analysis of the acute inflammatory response upon LPS injection in head kidney cells of *A. mexicanus*. (**a**) Experimental setup of the scRNAseq experiment using head kidney from surface fish (Río Choy) and cavefish (Pachón) 3hrs post injection of either PBS or an LPS mix in given concentration. After processing, cells were sequenced using 10X Genomics technology (see Methods for details), which resulted in given cell numbers and average genes per cell. (**b**) UMAP projection of PBS and LPS injected surface and cavefish. Given cell cluster are based on gene enrichment analysis for each cluster. See Data File 4 for gene enrichment in each cluster. (**c**) Heatmap of enriched genes within each cell cluster of control groups from surface fish and cavefish. Genes that were used for cell cluster identification are shown. For a complete heatmaps for PBS and LPS injected groups see Data Files 5-8.

Given the previously observed differences in the immune investment strategy between surface fish and cavefish, we speculated that these differences might be driven by genetic changes in hematopoietic stem cells (HSCs). While we found no changes in the relative abundance of HSCs across all samples (between 0.02 and 0.03, see Fig. 3b) we found distinct changes in the transcriptional profiles of HSCs between surface fish and cavefish control samples that may affect linage fate decision, the level of quiescence and self-renewal capacity (for a complete list of differential expressed genes in HSC see Data File 9). For example we found that cavefish cells show increased expression of *bcl3* (log2FC = 3.87), a marker for lymphoid progentitor cells^41^, and decreased expression of *npm1a* (log2FC = −1.22) and *mpx* (log2FC = −2.22), which are both key marker for early myeloid progenitor cells^42,43^. While further genetical analysis is needed to verify the genetic shift towards lymphoid progenitor cells in the cavefish HSCs, the transcriptional changes suggest that the different immune investment strategies in *A. mexicanus* could indeed be affected by genetic changes in cavefish HSCs. In addition, we found transcriptional changes that indicate differences in self-renewal capacity and quiescence of HSCs between surface fish and cavefish.

We detected increased expression of *c-myc* (*myca,* log2FC = 1.66) in cavefish cells, a positive regulator of HSC quiescence and self-renewal^44^. Furthermore, we found genes, such as *fscna* (log2FC = 3.056) and *pim1* (log2FC = 3.29), that are genetic marker for LT-(long-term) HSC significantly increased and genetic marker, such as *top2a* (log2FC = −4.72), *kif2c* (log2FC = −4.04) and *kif4* (log2FC = −3.05) for multipotent progenitor (MPP) cells significantly decreased in cavefish control samples compared to surface fish controls^45^. Interestingly, we found no expression of *cd34*, a common marker for mature HSCs and progenitor cells^46,47^, in any of the Pachón HSC cells, while we found expression of this marker in HSCs of the surface fish samples. Cd34 negative stem cells have been described as immature and quiescent with lower self-renewal capacity^47^. These findings point towards an increased proportion of immature LT-HSC in a more quiescent state in Pachón cavefish compared to surface fish.

Given the differences we observed in the lymphoid cell population structure between surface fish and cavefish, we next looked for differences in the LPS response of the lymphoid fraction. The expression of *interferon-γ* (*ifng*) of activated T-cells and Natural Killer (NK) cells is a well established response upon bacterial or viral infection. Although Pachón cavefish posses a high abundance of T-cells we found an increased abundance of *ifng* expressing cells in surface fish (Fig. 4a,b, relative abundance of *ifng* expressing cells in surface fish 0.041 vs. 0.016 in cavefish; for a complete list of differential expressed genes in the CD4+ T-cell cluster see Data File 10).

**Figure 4:**
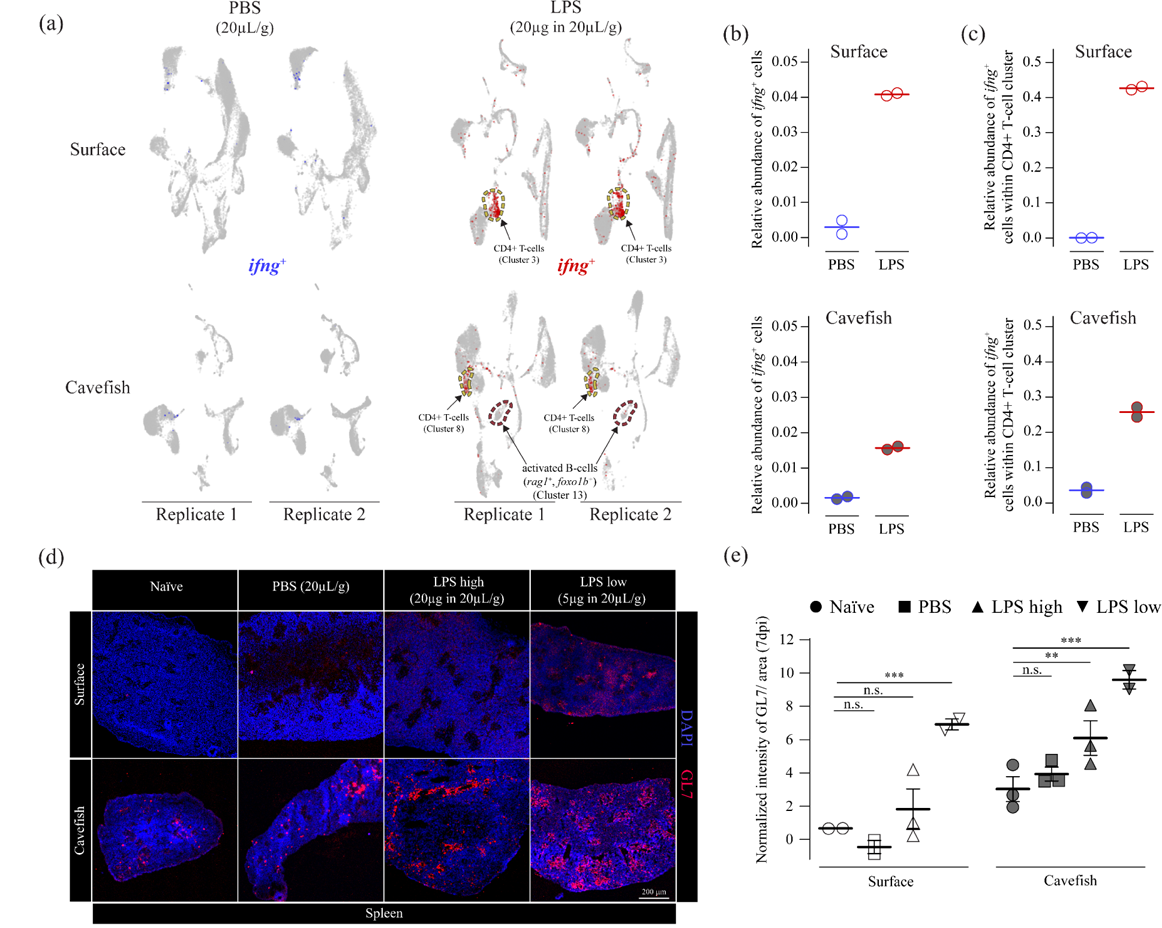
Adaptive response upon LPS injection in head kidney and spleen of surface and cavefish *A. mexicanus*. **(a)** *Interferon gamma* (*ifng*) expression 3hrs post PBS or LPS injection in head kidney cells. Upon LPS injection an activated B-cell cluster emerges in cavefish, which is absent in all other groups **(b)** Overall relative abundances for *ifng* expressing cells from scRNAseq experiment for each treatment group. **(c)** Relative abundances for *ifng* expressing cells from scRNAseq experiment within CD4+ T-cells for each treatment group. **(d)** Antibody staining (rat anti-GL7 Alexa Fluor® 647 (BD Pharmingen™) of activated B- and T-cells within germinal center of the spleen 7 days post injection (dpi) of either PBS (20µL/ g fish bodyweight), a high dose of LPS (LPS high; 20µg in 20µL/ g fish bodyweight) or a low dose of LPS (LPS low; 5µg in 20µL/ g fish bodyweight) or left naïve (not injected) as a control group. **(e)** Results of intensity analysis from (d) for 7 dpi. Images were analyzed from the 4 treatment groups from 2 timepoints (7 and 14 dpi, for 14 dpi analysis result see Figure S5) with n = 2-3 per treatment x population x timepoint sample. GL7 signal was quantified per area as defined by the DAPI signal using Fiji (see Methods for details) and intensities were normalized using the respective isotype control (see Methods for details). Empty symbols represent surface fish and filled symbols represent cavefish. A two-way ANOVA and subsequent multiple testing with FDR correction (Benjaminin-Hochberg) was used to detect differences between injected and control group (for complete statistical report see Data File 11). Significance values are indicated as ** for *p* ≤ 0.01 and *** for *p* ≤ 0.001.

While we did not detect a specific NK-cell cluster in any of the treatment groups, we observed that mainly CD4+ cells express *infg* in the acute pro-inflammatory response upon LPS injection. We also compared the relative abundance of *ifng* expressing cells within the CD4+ T-cell cluster and, again, found an increased abundance of *ifng* expressing cells in surface fish (Fig 4c, relative abundance of *ifng* expressing cells in the CD4+ T-cell cluster in surface fish 0.43 vs. 0.26 in cavefish). This decreased inflammatory response of CD4+ T-cells (Th1 response) in cavefish, suggests that cavefish lymphoid cells possess a different mode of response than surface fish upon bacterial recognition. Notably, we noticed a new activated B-cell population in LPS injected Pachón fish that is absent in Pachón PBS treated groups and in both surface fish treatment groups (Fig. 3b and 4a). The main charachteristic of this activated B-cell population is the expression of *foxo1b*, *rag-1* (Fig. 4a) and to a lesser extent *rag-2*, which is characteristic of activated B-cells^48,49^.

Based on this, we hypothesized that Pachón cavefish display increased activation of B-cells after injection with LPS compared to surface fish. To test this, we injected surface fish and Pachón cavefish with either PBS (20µL/g), a high dose of LPS (20µg in 20µL/g) or with a low dose of LPS (5µg in 20µL/g) or left naïve as control group and compared B- and T-cell activation status in the spleen, a major lymphoid organ for antigen processing in teleost fish^50^. To visualize activated B-cells we used an antibody against the GL-7 antigen that specifically stains activated B-cells that respond to a T-cell dependent antigen immunization event in the germinal center of lymphoid organs^51^. In surface fish we only found a significant increase of the GL-7 signal in the LPSlow group compared to naïve group at 7dpi (*p* ≤ 0.0001, multiple comparison after mixed effect analysis, Fig. 4d and e, see Data File 11 for detailed statistical analysis). For cavefish, however, we found significant increase of the GL-7 signal in the LPShigh and LPSlow group compared to the naïve group at 7dpi (*p* ≤ 0.01 and *p* ≤ 0.0001, respectively, Fig. 5e and d). We did not find a significant response upon LPS injection after 14dpi in surface and cavefish (see Fig. S5, see Data File 11 for detailed statistical analysis). These findings indicate that Pachón cavefish mount a more lymphoid (adaptive) driven immune response upon bacterial recognition than surface fish.

**Figure 5:**
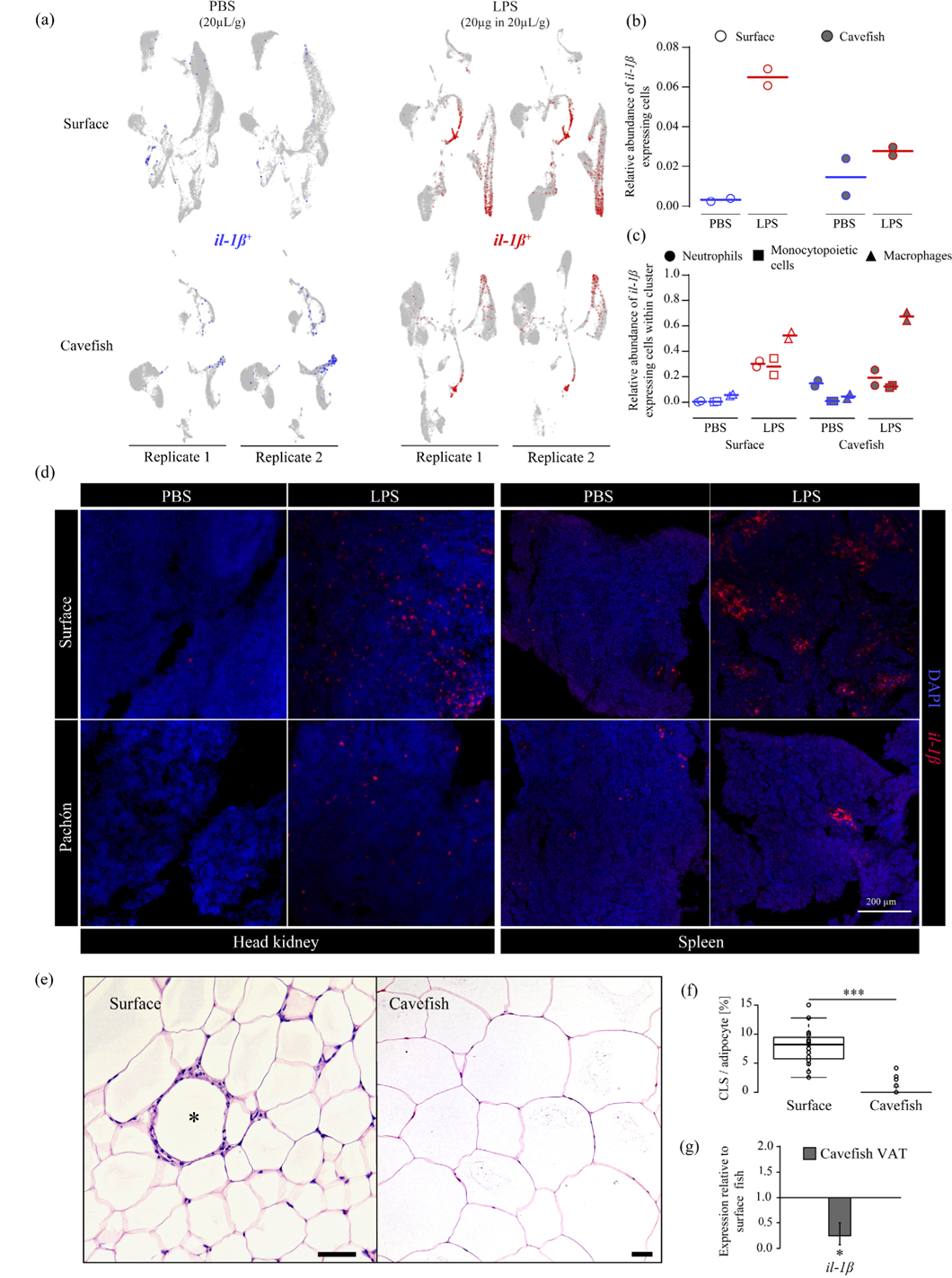
Reduced immune investment into myeloid cells alters inflammatory and immunopathological responses of *A. mexicanus*. *(a) Interleukin-1ß* (*il-1ß*) expression 3 hrs post PBS or LPS injection in head kidney cells. **(b)** Overall relative abundances of *il-1ß* expressing cells from scRNAseq experiment for each treatment group. **(c)** Relative abundances of *il-1ß* expressing cells from scRNAseq experiment within the main *il-1ß* expressing cell cluster for each treatment group. **(d)** *In-vivo* inflammatory response displayed by *in-situ* hybridyzation of *il-1β* using RNAscope in head kidney and spleen of surface fish and cavefish 3 hours post intraperinoteal injection of 20µg in µL/g(bodyweight) LPS. Images are representative of two independent experiments. Scale bar is 200 µm **(e)** H & E staining of visceral adipose tissue (VAT) of surface fish and cavefish. Crown-like structure (CLS) is indicated as * in surface VAT. Scale bar is 50 µm. **(f)** CLS count per 100 adipocytes in VAT of surface fish and cavefish in at least three fields of view for each fish (n=3). Significance values were determined by one-way ANOVA. For all box plots; center lines show the medians, crosses show means; box limits indicate the 25th and 75th percentiles as determined by R software; whiskers extend 1.5 times the interquartile range from the 25th and 75th percentiles, data points are represented by circles. Significance values are indicated *** for p ≤ 0.001. **(g)** Gene expression of *il-1β* in cavefish VAT relative to surface fish of the same fish that were used for (f). Significance values were determined by a pairwise fixed reallocation randomization test using REST2009 software and are indicated as * for p ≤ 0.05.

Finally, we asked whether the reduced investment into innate immune cells, such as granulocytes and monocytes, in cavefish is accompanied by changes in gene expression of the proinflammatory response (for a complete list of differential expressed genes in the neutrophil and macrophage cluster see Data File 12 and 13, respectively). Here we found that the main innate immune cells, neutrophils and macrophages, from cavefish showed increased expression of *csf3r* (log2Fold = 2,85 and 1,14 compared to the surface fish PBS group, respectively). Although we also found increased expression of the macrophage-associated colony stimulating factor receptor (*csf1r*) in the cavefish control group compared to surface control fish, expression was very low in all groups. However, the increased expression of *csf3r* in neutrophils and macrophages indicates a higher sensitivity for its ligand *csf3* (*g-csf*), which is produced by a variety of different immune cells to stimulate the release of Csf3r positive cells into the bloodstream. We also found distinct changes in the inflammatory response in neutrophils and macrophages of surface fish and cavefish upon injection with LPS. For example, components of the NF-κB pathway, a major regulator of inflammatory processes in vertebrates^52^, was significantly increased in surface fish. However, based on the overall reduced investment of cavefish into innate immune cells we asked whether there is also a reduction of cells that mediate pro-inflammatory responses.

To test this, we used the expression of the cytokine *il-1β* as a readout, as this cytokine is described as one major regulator of proinflammatory respones in teleost fish^53^. We detected induced expression in granulopoietic cells (mainly mature neutrophils), monocytopoietic cells (mainly mature monocytes) and macrophages in surface fish and cavefish upon LPS injection (Fig. 5a). We found a 2.3 – fold increase of *il-1β* expressing cells in surface fish 3 hrs post LPS injection compared to cavefish (mean overall relative abundance of cells expressing *il-1β* in surface fish 0.065 vs. 0.028 in cavefish, see Fig. 5b). Interestingly, cavefish seemed highly variable after PBS injection (Fig. 5b), but when we compared expression of *il-1β* within each cell cluster, only cavefish neutrophils showed elevated *il-1β* expression in both replicates of the PBS injected group that is comparable to the induced expression after LPS injection (Fig. 5c). This, however, is a cavefish neutrophil specific phenomenon since monocytopoietic cells (containing mature monocytes) and macrophages do not express *il-1β* in the PBS injected Pachón samples (Fig. 5c). It is noteworthy that macrophages from surface fish and cavefish showed the highest increase in *il-1β* expressing cells and represent presumably the main producer of *il-1β* upon LPS injection in A. *mexicanus* (Fig. 5c).

To verify the reduction in neutrophils and monocytes / macrophages that can initiate a proinflammatory response, we designed an *in-situ* RNAscope probe for *il-1β* to visualize *il-1β* expression in head kidney and spleen. We injected surface fish (Río Choy) and cavefish (Pachón) with 20µg in 20µL/ g LPS and dissected head kidney and spleen 3 hours post injection (see Methods section for details). In line with the scRNAseq analysis, LPS injected surface fish showed an increased number of *il-1β* positive cells in the head kidney compared to cavefish (Fig. 5d). We also detected considerably fewer cells that express *il-1β* after injection with LPS in the spleen from cavefish compared to surface fish (Fig. 5d). In teleost fish, the spleen contains high numbers of mononuclear phagocytes, e.g. macrophages^32,54^ but is generally not a hematopoietic tissue for such cell types ^50^. In addition, we also used the *il-1β* RNAscope probe on dissociated head kidney cells from surface fish 3 hrs post injection with LPS and we were able to validate that mainly cells with monocytic and neutrophilic characteristics (multi-lobbed nuclei) express *il-1β* (Fig. S6).

Based on these results, we hypothesized that the lack of cells that initiate a systemic pro-inflammatory response in cavefish upon exposure to an immune stimulant (e.g., LPS) could potentially lead to a decreased presence of immunopathological phenotypes that result from such pro-inflammatory responses. Cavefish produce substantially more visceral adipose tissue (VAT) than surface fish^26^. In mammals, the amount of VAT is positively correlated with number of monocytes infiltrating the adipose tissue and mediating inflammatory processes resulting in the formation of crown-like structures (CLS)^55^. Therefore, we tested whether VAT of *A. mexicanus* shows signs of CLS and if surface fish and cavefish differ in their occurrence. We detected CLS in the visceral adipose tissue of surface fish (Fig. 5e), but not in cavefish, despite the prevalence of large, hypertrophic adipocytes (average numbers of CLS in 100 adipocytes were 7.95 for surface vs. 0.6 for cavefish, *p* ≤ 0.001, one-way ANOVA, Fig. 5f). To measure levels of *il-1β* expression, we took a sub-sample of the VAT for RT-qPCR analysis. We detected reduced expression of *il-1β* in VAT of cavefish relatively to surface fish (mean relative expression of cavefish compared with surface fish 0.249, *p* ≤ 0.05, pairwise fixed reallocation randomization test, Fig. 5g). In combination with the reduced number of crown-like structures, our data indicate a reduction of pro-inflammatory granulocytes and macrophages in VAT of cavefish potentially enabling increased VAT storage in cavefish without immunopathological consequences.

## Conclusion

Our study elucidates how adaptation to low biodiversity in caves affects the immune investment strategy of a vertebrate host. Besides differences in biodiversity, there are a variety of environmental parameters that differ between the river and cave habitat (e.g. light conditions, food availability, oxygen concentration). Differences in these parameters may potentially influence different physiological systems that affect immune cell composition and function of *A. mexicanus*. However, given that the maintenance and the control of the immune system is costly too^2^, the changes in the immune investment strategy of cavefish is likely an evolutionary response facilitated by the low parasite diversity in the cave environment. Proinflammatory reactions are one of the main causes for immunopathological phenotypes and have a tremendous impact on the fitness of an organism and can be caused by a variety of environmental factors^56-58^. We interpreted the reduction of innate immune cells in cavefish, which mediate proinflammatory processes and act against parasites, as an adaptation decreasing auto-agressive immunopathology from a hyper-sensitive immune system in an environment without parasite diversity. With *A. mexicanus* we present a vertebrate system, which lost parasite diversity for thousands of generations and presents immunological adaptations to such an environment that prevent immunopathology.

## Supporting information

Data File 1_Statistics_Phagocytosis

Data File 2_Cell Gallery Cluster

Data File 3_Cluster specific gene expression_Figure2

Data File 4_Cluster specific gene expression_Figure3

Data File 5_SurfacePBSMarkerHeatmap

Data File 6_SurfaceLPSMarkerHeatmap

Data File 7_CavefishPBSMarkerHeatmap

Data File 8_CavefishLPSMarkerHeatmap

Data File 9_HSC_Cavefish_vs_Surface_PBS

Data File 10_Cd4T-cell_Cavefish_vs_Surface_PBS and LPS

Data File 11_StatiticsGL7Intensity

Data File 12_Neutrophils_Cavefish_vs_Surface_PBS and LPS

Data File 13_Macrophages_Cavefish_vs_Surface_PBS and LPS

## Acknowledgements

We are grateful to the cavefish facility staff at the Stowers Institute for support and husbandry of the fish. We would like to thank the staff from the Histology core at the Stowers Institute for their technical support, Jillian Blanck from the Cytometry core for performing the sorting of head kidney cells, Michael Peterson, Allison Peak and Anoja Perera for the scRNA-seq support, Mark Miller for his support on the fish anatomy figure and Hua Li for her support on the statistical analysis. Furthermore, we would like to thank Sean A. McKinney for providing the ImageJ macro for GL7 quantification. The authors also kindly acknowledge Joachim Kurtz for helpful discussions. NR was supported by institutional funding, the Edward Mallinckrodt foundation and the JDRF. RP was supported by a grant from the Deutsche Forschungsgemeinschaft (PE 2807/1-1).

## Author Contributions

RP and NR conceived of the study. RP designed and coordinated the experiments with support from ACB and JK. RP, JLP, AK and EM collected, dissected and examined cave and surface wild populations with support from JPS. RP performed and analysed immune assays, flow cytometry experiments and histological analysis with support from ACB, YW, DT and BPS. SC performed single cell sequencing analysis with support from RP. RNAscope experiments and analysis were performed by YW, DT and BPS with support from RP and JK. RP and NR designed and RP made the figures. RP and NR wrote the paper and all authors read and edited the paper.

### Data availability statement

Original data underlying this manuscript can be accessed from the Stowers Original Data Repository at http://www.stowers.org/research/publications/libpb-1391. The scRNA-seq data generated by Cell Ranger can be retrieved from the GEO database with accession number GSE128306.

## Methods section

### Field sample collection

Collection for this study was conducted under permit No. SGPA/DGVS/03634/19 granted by the Secretaría de Medio Ambiente y Recursos Naturales to Ernesto Maldonado. Study sites are located in the Sierra de El Abra region of northeastern Mexico in the states of San Luis Potosí and Tamaulipas. The El Abra region experienced repeated uplift and erosional events that carved the underground limestone caverns (Mitchell, Russell, & Elliott, 1977). We collected samples from Pachón cave; one of 30 caves in the region with known cave-dwelling *Astyanax* populations (Espinasa et al., 2018; Mitchell et al., 1977). We also collected samples of the surface morphotype from Nacimiento Río Choy approximately 95km south of Pachón cave.

We collected fish from Pachón cave and Nacimiento Río Choy on July 12th, 13th and 14th 2019 during the rainy season. Pachón fish were collected in the morning of July 12th using handheld nets. Río Choy fish were collected during the day on July 13th and 14th using a combination of handheld nets, net traps and a modified plastic bottle trap. Captured fish were placed in their environmental water and euthanized on the day of capture.

### Fish husbandry

Unless otherwise stated, all fish used for the experiments were adult female fish aged 12-16 months. Surface morphs of *Astyanax mexicanus* were reared from Mexican surface fish (Río Choy) and cavefish originated from the Pachón and Tinaja cave. Fish were housed at a density of ∼ 2 fish per liter. The aquatic animal program at the Stowers Institute meets all federal regulations and has been fully AAALAC-accredited since 2005. *Astyanax* are housed in glass fish tanks on racks (Pentair, Apopka, FL) with a 14:10 h light:dark photoperiod. Each rack uses an independent recirculating aquaculture system with mechanical, chemical and biologic filtration and UV disinfection. Water (supplemented with Instant Ocean Sea Salt [Blacksburg, VA]) quality parameters are maintained within safe limits (Upper limit of total ammonia nitrogen range, 1 mg/L; upper limit of nitrite range, 0.5 mg/L; upper limit of nitrate range, 60 mg/L; temperature, 22 °C; pH, 7.65; specific conductance, 800 μS/cm; dissolved oxygen 100 %). Fish were fed once per day with mysis shrimp and twice per day with Gemma diet (according to the manufacturer is Protein 59%; Lipids 14%; Fiber 0.2%; Ash 14%; Phosphorus 1.3%; Calcium 1.5%; Sodium 0.7%; Vitamin A 23000 IU/kg; Vitamin D3 2800 IU/kg; Vitamin C 1000 mg/kg; Vitamin E 400 mg/kg) at a designated amount of approximately 3% body mass. Routine tank side health examinations of all fish were conducted by dedicated aquatics staff twice daily. *Astyanax* colonies are screened at least biannually for *Edwardsiella ictaluri, Mycobacterium spp., Myxidium streisingeri, Pseudocapillaria tomentosa, Pseudoloma neurophilia*, ectoparasites and endoparasites. At the time of the study, none of the listed pathogens were detected.

### *In-vitro* gene expression analysis

Single cell suspensions from freshly dissected head kidney tissue were produced by forcing tissue through 40 µM cell strainer into L-15 media (Sigma), containing 10 % water and 5 mM HEPES buffer (pH 7.2) and 20 U/mL heparin (L-90). The strainer was washed once with L-90 and cells were washed once by spinning cells at 500 x *g* at 4 °C for 5 mins. Supernatant was discarded and cells were resuspended in 1 mL of L-90 media (L-15 containing 10 % water, 5 mM HEPES (7.2 pH), 5 % fetal calf serum, 4 mM L-glutamin, Penicillin-Streptomycin mix with 10,000 U/mL each). Cells were counted using EC800 analyser (Sony Biotechnology) and 1×10^6^ cells were plated in 48 well plate in 500 µL and incubated over night at 21 °C. At timepoint 0, 20 µg / ml or 0.2 µg / mL lipopolysaccharide mix in PBS (*Escherichia coli* O55:B5 and *E. coli* O111:B4, 1 mg/mL each) or PBS alone as a control was added to the cells, respectively. After 1, 3, 6, 12 and 24 hours, cells were harvested and immediately snap frozen in liquid nitrogen and RNA was isolated as described previously^59^. 100 ng of RNA (concentration was measured using the Qubit system (Thermo Fisher)) from each sample was used for cDNA synthesis using the SuperScript™ III First-Strand Synthesis System kit (Invitrogen) following manufacturer instructions. Resulting cDNA was used for RTqPCR using the PerfeCTa® SYBR® Green FastMix® (Low ROX) (Quantabio) following manufacturer instructions. Gene specific primers (see Table S1) were used for amplification of target and the two housekeeping genes (rpl32 and rpl13a, see Table S1 for details). Where possible, gene specific primers were designed to span an exon – exon junction. Samples were pipetted in a 384 well plate using a Tecan EVO PCR Workstation (Tecan) and samples were run in technical triplicates on a QuantStudio 7 Flex Real-Time PCR System (Thermo Fisher). Quality control for each sample was performed using the QuantStudio Real-Time PCR software (Thermo Fisher) and data was exported for analysis in REST 2009^60^ as described before^59^. PBS control samples from each time point and sample was used as the reference to calculate relative expression of target genes for each timepoint and fish, respectively.

### Phagocytosis Assay

Phagocytosis was measured as previously described^61^. Briefly, a single cell solution from freshly dissected head kidney tissue was prepared as described above and 4×10^5^ cells were pipetted into 96 well flat bottom plate and Alexa-488 tagged *Staphylococcus aureus* (Thermo Fisher) were added in a 1:50 cells / bacteria ratio. To control for cell viability a sample without bacteria was included and to control for active phagocytosis a sample with cells containing bacteria and cytochalasin B (CCB) (0.08 mg /mL) for each individual sample was included. Cells were incubated in 200 µL of L-90 media at 21 °C for 1 and 3 hours, respectively. To exclude dead cells and signal from non-phagocytosed particles, cells were stained with Hoechst and all samples were quenched using 50 µL Trypan Blue (0.4 % solution, Sigma) before the measurement. Samples were measured on EC800 Analyzer (Sony Biotechnology). Cells were gated for live and Alexa-488 positive and phagocytosis rate was calculated as the ratio of live (Hoechst positive, Excitation 352 nm, Emission 461 nm, FL-6) and phagocytes (Alexa-488 positive, Excitation 495 nm, Emission 519 nm, FL-1) vs. live cells.

### Scatter analysis of head kidney

Single cells from head kidney from adult surface fish were extracted as described above. Cells were stained with DAPI to exclude dead cells and live cells were sorted based on populations as described in Fig. 1i using forward side scatter and side scatter characteristics of cells using an Influx System (BD). 1000 cells per population were sorted on a Thermo Scientific™ Shandon™ Polysine Slides and incubated for 30 min at 21 °C, so cells could settle and adhere to slides. Cells were then fixed with 4 % Paraformaldehyde and washed three times in PBS. Cells were then stained using May-Grünwald Giemsa protocol. Briefly, slides were stained for 10 min with a 1:2 solution of May-Grünwald (made in phosphate buffer pH 6.5, filtered), the excess stain was drained off and slides were stained 40 min with a 1:10 solution of Giemsa (made in phosphate buffer pH 6.5, filtered). Then slides were rinsed in ddH_2_0 by passing each slide under running ddH_2_0 10 times. For differentiation, a drop of 0.05 acid water (5ml glacial acetic acid/95ml ddH_2_O) was put on slide for approx. 4 seconds and quickly rinsed off. Slides were rinsed well, air dried and coverslipped. Only cells that could clearly be identified based on studies in closely related organisms using a similar approach^30,32^ were used as representative image for Fig. 1i.

### Image-based cluster analysis of head kidney

For this analysis, hematopoietic cells from the head kidney were presorted to remove the mature erythrocyte cluster using the S3 Cell Sorter (Bio-Rad) using scatter features (as in Fig. 1i). This was necessary since mature erythrocytes account for about 8 % and 7 % respectively in surface fish and Pachón fish of the entire cell count in the head kidney based on the erythrocyte population we were able to identify using scatter alone (see Fig. 1i). However, based on their biconcave morphology we found erythrocytes in all populations that we could separate through scatter, although mainly in the myelomonocytic and progenitor populations to different degrees since different orientations in the flow cell of erythrocytes result in different morphological shapes^37^. Based on this, the presence of mature erythrocytes results in massive over-clustering^37^. Reduction of erythrocytes through sorting based on scatter can be used to reduce the amount of over-clustering using the pipeline^37^. Sorted cells were stained with 5 µM Draq5 and 10,000 nucleated, single events were acquired from samples on the ImageStream®X Mark II at 60x, slow flow speed, using 633 nm laser excitation. Bright field was acquired on channels 1 and 9 and Draq5 on channel 11. SSC was acquired on channel 6. Intensities from 25 unique morphological features were extracted. Further analysis was done as described before^37^.

### Intraperitoneal injection of LPS

Fish were individualized and fasted the day before the treatment (naïve, PBS-injected or LPS injected). After 24hrs the fish were anesthetized using ice cold system water and either PBS (20µL/g bodyweight) or a LPS mix (*E. coli* O55:B5 and *E. coli* O111:B4, 20µg or 5µg in 20 µL/g bodyweight) was injected intraperitoneally using an insulin syringe (3/10 mL, 8 mm length, gauge size 31G, BD). After given timepoints, fish were euthanized using buffered Tricaine solution (500 mg/L) and respective organs were dissected and were either dissociated (single cell RNA sequencing) or immediately fixed in 4 % paraformaldehyde / DEPC water (RNAscope analysis) for subsequent analysis.

### Single cell RNAseq

Dissociated hematopoietic cells were stained with DAPI to exclude dead cells and live cells were sorted based on populations as described in Fig. 1i, where only myelomonocyte, lymphocyte and progenitor populations were sorted in L-90 media, to reduce the relative abundance of mature erythrocytes. Sorted cells were spun down (500 x rcf, 4° C, 5 min), supernatant was discarded, and cells were resuspended in L-90 media and run again on Influx system to ensure removal of mature erythrocyte cluster and measure cell viability (percentage live cells after sorting: surface fish 82.2 % and cavefish 88.9 %). Cells were loaded on a Chromium Single Cell Controller (10x Genomics, Pleasanton, CA), based on live cell concentration, with a target of 6,000 cells per sample. Libraries were prepared using the Chromium Single Cell 3’ Library & Gel Bead Kit v2 (10x Genomics) according to manufacturer’s directions. Resulting short fragment libraries were checked for quality and quantity using an Agilent 2100 Bioanalyzer and Invitrogen Qubit Fluorometer. Libraries were sequenced individually to a depth of ∼330M reads each on an Illumina HiSeq 2500 instrument using Rapid SBS v2 chemistry with the following paired read lengths: 26 bp Read1, 8 bp I7 Index and 98 bp Read2. Raw sequencing data were processed using 10X Genomics Cell Ranger pipeline (version 2.1.1). Reads were demultiplexed into Fastq file format using cellranger mkfastq. Genome index was built by cellranger mkref using cavefish genome astMex1, ensembl 87 gene model. Data were aligned by STAR aligner and cell counts tables were generated using cellranger count function with default parameters. Cells with at least 500 UMI counts were loaded into R package Seurat (version 2.3.4) for clustering and trajectory analysis. 4991 cells for surface and 4103 cells for Pachón cavefish were used for downstream analysis. The UMI count matrix were log normalized to find variable genes. First 12 principal components were selected for dimension reduction and t-SNE plots. Marker genes were used to classify clusters into lymphocytes, myelomonocytes and progenitor types. The results generated by Cell Ranger can be retrieved from the GEO database with accession number GSE128306. The assignment of cell identities is based on their transcription profile determined by similar approaches in zebrafish^32,62-65^.

### Single-cell RNAseq for LPS injection

Libraries were sequenced paired-end using Illumina Novaseq S2 flowcell. Raw data were processed using Cell Ranger pipeline (version 3.0) and demultiplexed into Fastq file format using cellranger mkfastq. Data were aligned to astMex1, ensemble 87 gene model by STAR aligner and cell counts tables were generated using cellranger count function with default parameters. Cells with at least 500 UMI counts were loaded into R package Seurat (version 3.0) for clustering and trajectory analysis. The UMI count matrix were log normalized to find top 2000 variable genes using vst selection method from Seurat. Replicates were then integrated using SCTtransform function FindIntegrationAnchors based on Seurat’s vignettes. Principal components cutoffs were selected based on Jackstraw and Elbowplot function for dimension reduction and UMAP plots. De novo markers genes were generated using FindAllMarkers function and to plot heatmaps in Figure 3. Trajectory analysis were computed using R package slingshot. All scRNA-seq data can be retrieved from the GEO database with accession number GSE128306.

### *Il-1β* RNAscope assay

Section preparation and RNA in situ hybridization were performed as previously reported^26,66^. Briefly, for tissue section, respective tissues (head kidney, spleen) were dissected from surface fish and cavefish, followed by immediate immersion into 4% PFA in DEPC H_2_O (diluted from 16% (wt/vol) aqueous solution, Electron Microscopy Sciences, cat# 15710) for 24hr at 4°C to fix the tissue, then rinsed well with 1xPBS, dehydrated through graded ethanol (30%, 50%, 70%) and processed with a PATHOS Delta hybrid tissue processor (Milestone Medical Technologies, Inc, MI). Paraffin sections with 8 µm thickness were cut using a Leica RM2255 microtome (Leica Biosystems Inc. Buffalo Grove, IL) and mounted on Superfrost Plus microscope slides (cat# 12-550-15, Thermo Fisher Scientific). For single cell solutions, head kidney and spleen were dissected, and single cell solutions were produced as described above and approx. 20 µL of the suspension was pipetted on Superfrost Plus microscope slides (cat# 12-550-15, Thermo Fisher Scientific). Cells were allowed to settle for 30 min and fixed using 4% PFA (diluted from 16% (wt/vol) aqueous solution, Electron Microscopy Sciences, cat# 15710) for 1hr at RT, then rinsed well with 1XPBS, dehydrated through graded ethanol (30%, 50%, 70%). RNA in situ hybridization was performed using RNAscope multiplex fluorescent detection V2 kit according to the manufacturer’s instructions (Advanced Cell Diagnostics, Newark, CA). RNAscope probe for *il-1β* was a 16ZZ probe named Ame-LOC103026214-C2 targeting 217-953 of XM_022680751.1.

Images of sections were acquired on a Nikon 3PO spinning disc on a Nikon Ti Eclipse base, outfitted with a W1 disk. A 0.75 NA, Plan Apochromat Lambda 20x air objective was used. DAPI and AF647 were excited with a 405 nm and 640 nm laser, respectively, with a 405/488/561/640 nm main dichroic. Emission was collected onto an ORCA-Flash 4.0 V2 digital sCMOS camera, through a 700/75 nm and 455/50 nm filter for the far-red channel and DAPI channel, respectively. Z-step spacing was 1.5 microns. All microscope parameters and acquisition were controlled with Nikon Elements software. Identical camera exposure time and laser power was used across samples. All image processing was done with an open source version of FIJI^67^ with standard commands. A Gaussian blur with radius of 1 was applied and a rolling ball background subtraction with a radius of 200 pixels was applied to every channel with the exception of the DAPI channel. Following that, a max projection across the slice was applied. For direct comparison, images shown are contrasted identically in the far red channel (*il-1β*).

### GL-7 analysis of spleen after LPS injection

Fish were dissected 3hrs after treatment and the spleen was immediately embedded with OCT compound (Tissue-Tek, CA) and freezed at −70°C. Cryo sections with 12µm thickness were cut using a Leica CM3050S cryostat (Leica Biosystems Inc. Buffalo Grove, IL) and mounted on glass slides. Sections were kept in cyrostat for 2hr before fixed with pre-chilled 75% acteone/25% ethynoal at room temprature for 30 min. Immunofluorescence assay was performed manually using an Alexa Fluor 647 conjugated rat anti-mouse T-cell and B-cell activation antigen (BD Pharmingen, GL7 clone, cat# 561529) and a matched isotype control (BD Pharmingen, R4-22 clone, cat# 560892). Here, IF experiments were repeated three times with different populations and timepoints, which accumulated a total number of 48 animals. In brief, sections were rehydrated with 1X PBS and background was blocked by incubating sections in Background Buster solution (NB306, Innovex Biosciences, CA, USA) for 30 min. The antibody was diluted 1:500 in Antibody Diluting Reagent (003118, Invitrogen, Carlsbad, CA, USA) and incubated overnight at 4°C. Sections were further stained with DAPI (1:1000) for 10min, and then washed in tris-buffered saline (25 mM Tris, 0.15 mM NaCl, pH7.2) with 0.05% Tween-20 (TBST) and coverslipped before imaging. Images of sections were acquired on a Zeiss LSM 700 upright microscope. A 5x air objective was used. DAPI and AF647 were excited with a 405 nm and 640 nm laser, respectively, with a 405/488/561/640 nm main dichroic. Emission was collected onto an ORCA-Flash 4.0 V2 digital sCMOS camera, through a 700/75 nm and 455/50 nm filter for the far-red channel and DAPI channel, respectively. All image processing was done with an open source version of FIJI^67^ with standard commands. For GL-7 intensity analyis we used Fiji macro that is publicly available under https://github.com/jouyun/smc-macros/blob/master/ROP_IntensityMeasurement.ijm.

### Visceral adipose tissue analysis

We dissected the visceral adipose tissue (VAT) from the abdominal cavity as described previously^26^. In short, we manually removed the intestinal sack of the fish and carefully isolated a piece of fat tissue located around the gut for RTqPCR as described above. The rest of the sample was immediately fixed in 4% paraformaldehyde for 18 h at 4 °C and embedded in JB-4 Embedding solution (Electron Microscopy Sciences; #14270-00) while following kit instructions for dehydration, infiltration and embedding. After sectioning at 5 μm, we dried slides for 1 h in a 60°C oven and stained slides with hematoxylin for 40 min. After rinsing the slides in PBS, semi-dried slides were stained with eosin (3% made in desalted water) for 3 min. Slides were washed with desalted water and air dried. At least 3 images from VAT of each fish were taken at similar location around the gut and crown-like structures were scored as described previously^68^. Images were obtained using a 10X objective on Zeiss Axioplan2 upright microscope and adipocytes and CLS were counted using Adobe Photoshop CC (Version 19.1.0).

### Statistical Analysis

Graphical data and statistics were produced using R^69^ except otherwise stated. For comparisons between populations we used a one-way ANOVA and corrected for multiple testing against the same control group (FDR) with Benjamini-Hochberg test^70^. For analysis of RT-qPCR data we used the REST2009 software where significant differences between two groups were determined by a pairwise fixed reallocation randomization test^60^. Two-way ANOVA analysis was done using Graph Pad Prism Software (Version 8.0.2). Multiple testing against the same control group was corrected with FDR test Benjamini-Hochberg test^70^. To determine significant differences of morphological cell cluster between surface fish and cavefish that resulted from X-shift clustering^71^ we used a negative binominal regression model as described before^37^.

### Animal experiment statement

Research and animal care were approved by the Institutional Animal Care and Use Committee (IACUC) of the Stowers Institute for Medical Research.

## Supplemental Figures

**Figure S1:**
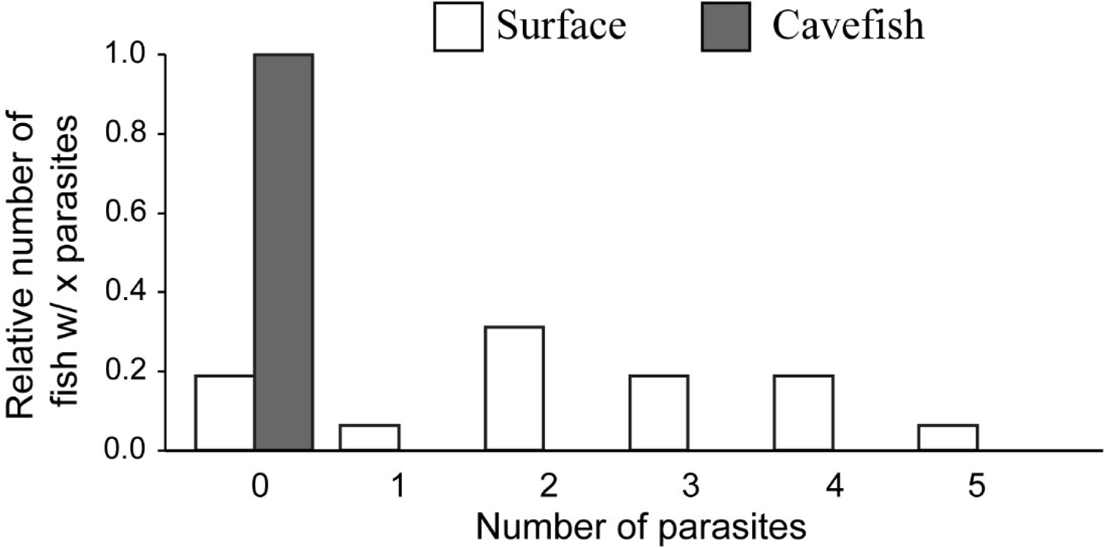
Numbers of different macroparasites found in wild surface (Río Choy) or cavefish (Pachón).

**Figure S2:**
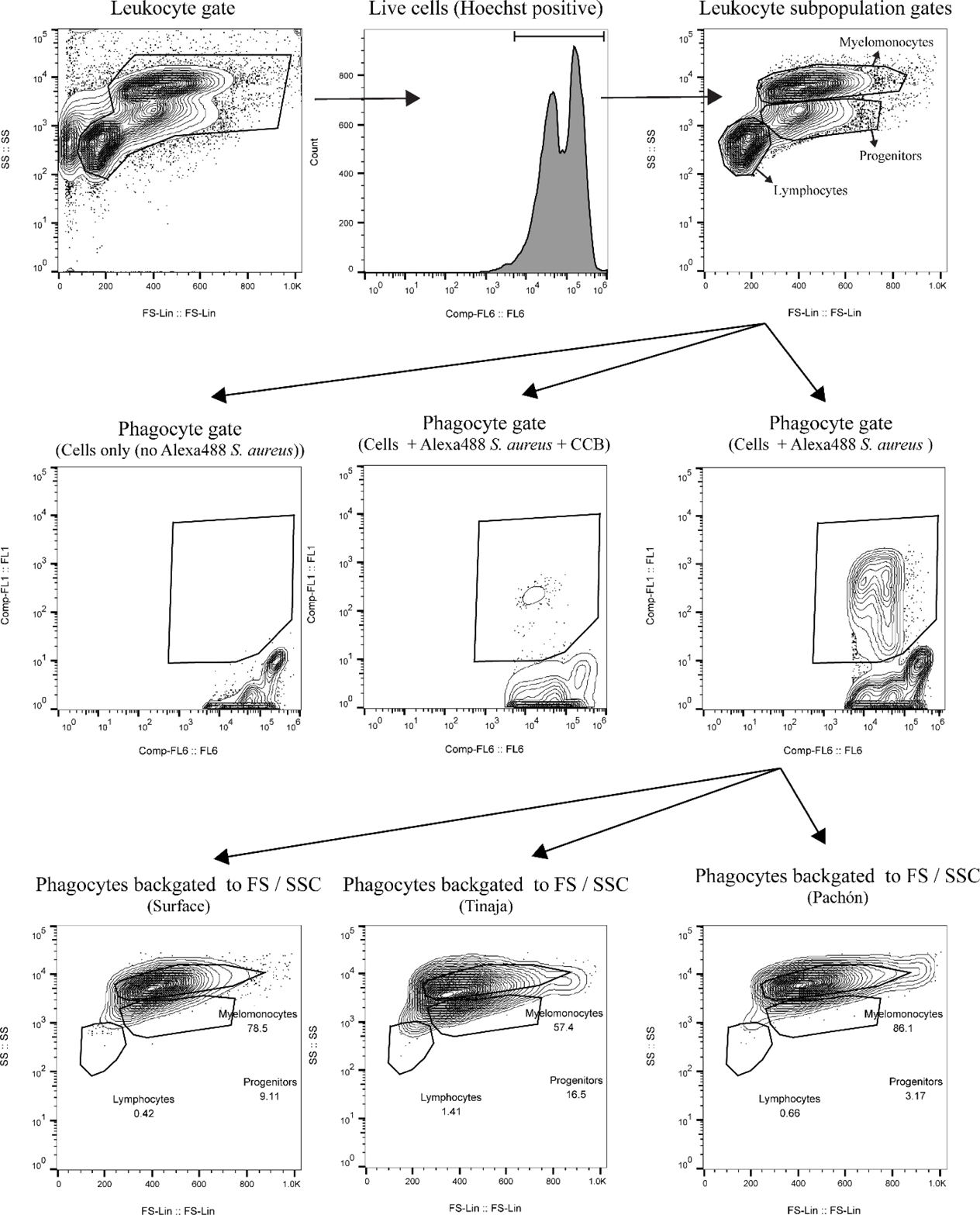
Gating strategy to identify phagocytes in head kidney cell suspension from Astyanax mexicanus using Alexa-488 Staphylococcus aureus and cytochalasin-B (CCB) as a control for active phagocytosis.

**Figure S3:**
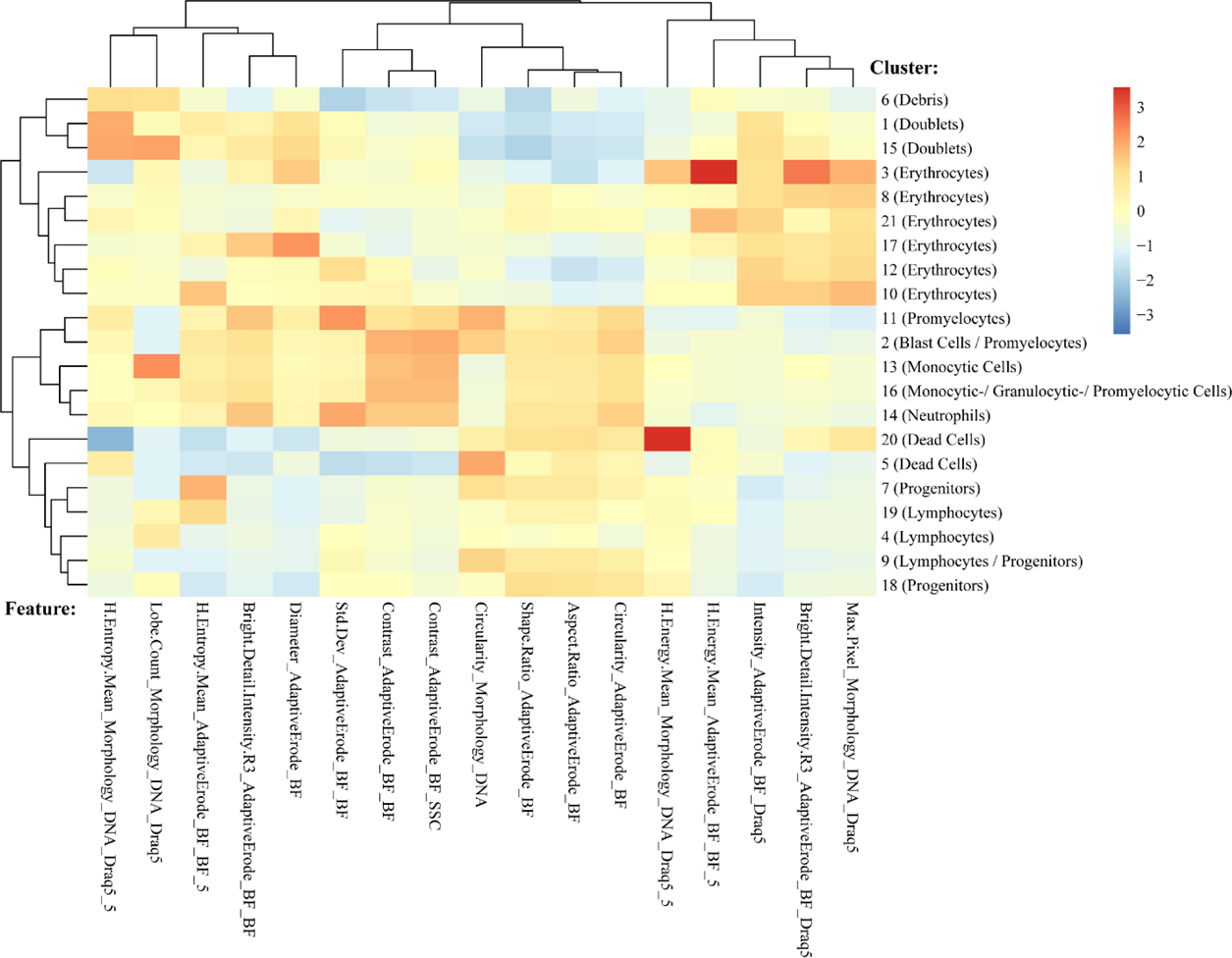
Spearmen Correlation of overall mean feature intensities used for clustering and single cluster after X-shift based clustering using head kidney cells from surface fish and cavefish. See Table S3 for details of features.

**Figure S4:**
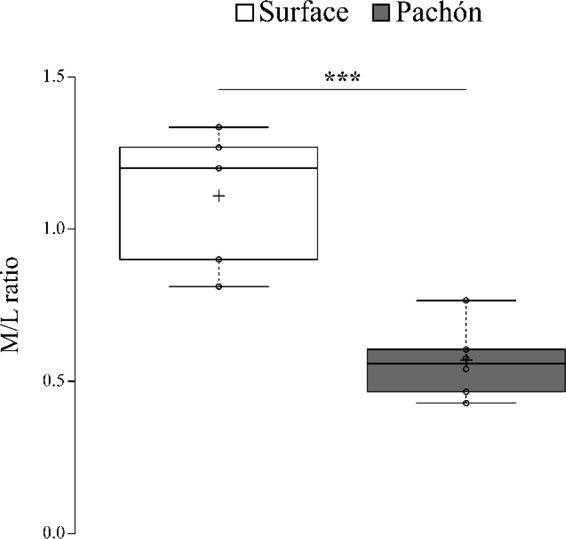
Myelomonocyte/lymphocyte ratio based on Image3C analysis. Differences between surface fish (Río Choy) and cavefish (Pachón) were tested using a one-way Anova. Significances are indicated as *** for p ≤ 0.001.

**Figure S5:**
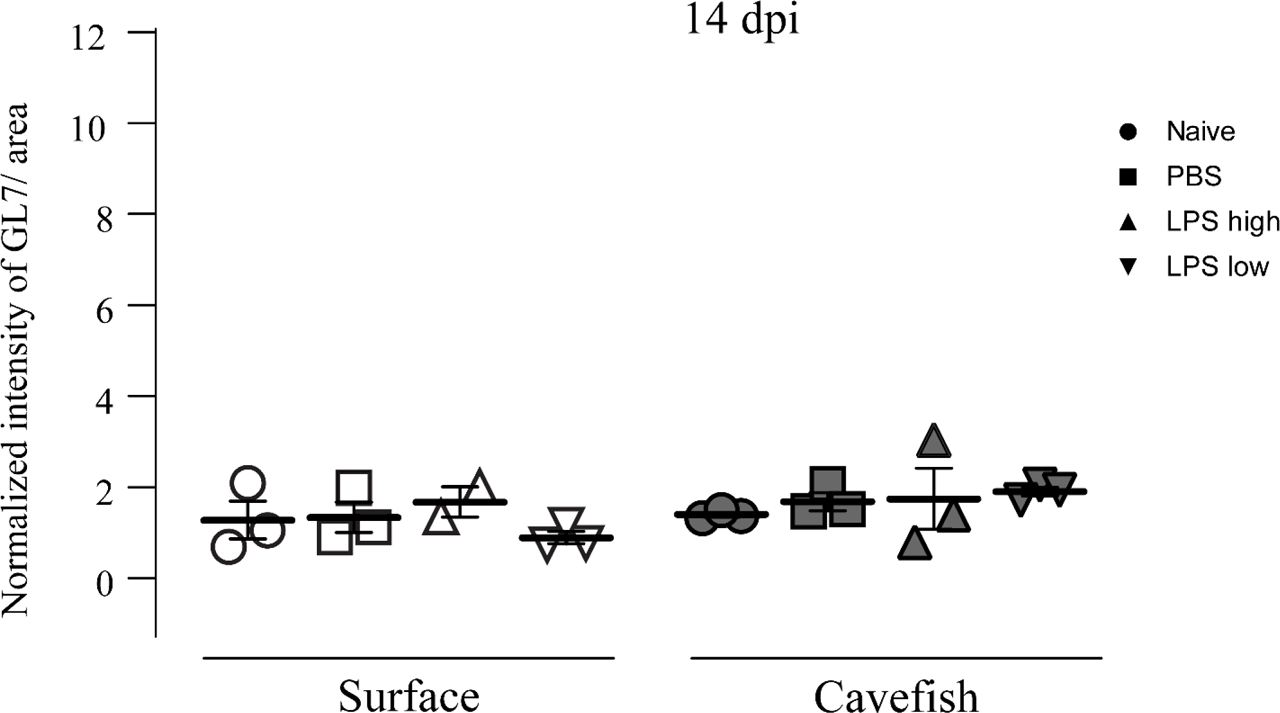
Antibody staining (rat anti-GL7 Alexa Fluor® 647 (BD Pharmingen™) of activated B- and T-cells within germinal center of the spleen 14 days post injection (dpi) of either PBS (20µL/g fish bodyweight), a high dose of LPS (LPS high; 20µg in 20µL/ g fish bodyweight) or a low dose of LPS (LPS low; 5µg in 20µL/ g fish bodyweight) or left naïve (not injected) as a control group.

**Figure S6:**
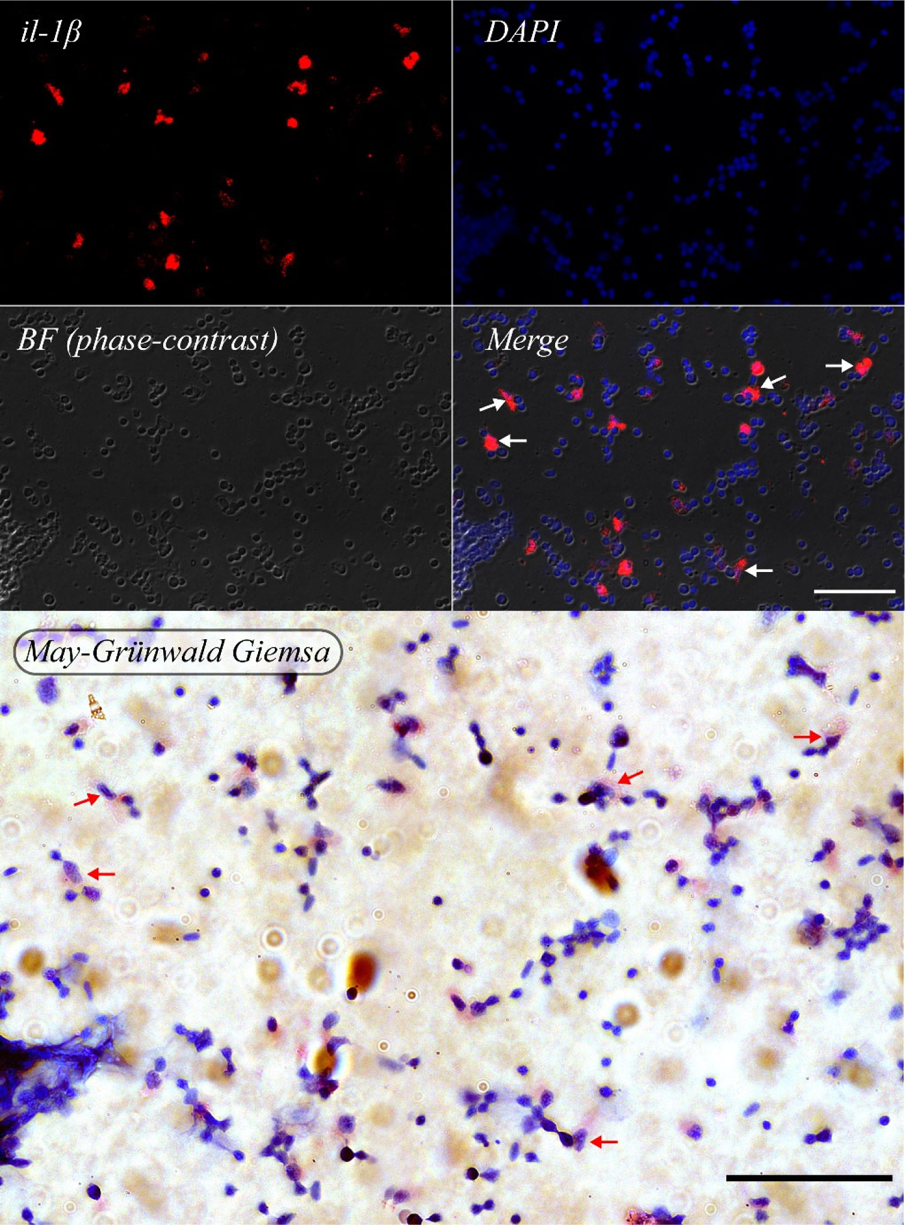
RNAscope analysis on dissociated head kidney cells from surface fish 3hpi with LPS (20µg in 20µL/g bodyweight). Dissociated cells were transferred on Microscope slide and stained as described in Supplemental Methods. After RNAscope imagining using 700/75 nm and 455/50 nm filter for the far red channel (*il-1β*) and DAPI channel, respectively, cover slip was removed and cells were stained after May-Grünwald Giemsa. Exact position of slide was imaged with 63 X Oil objective as before. Cells expressing *il-1β* are marked with white arrow in ‘*Merge*’ image and the same cells are marked with a red arrow in ‘*May-Grünwald Giemsa*’ image.

**Table S1:**
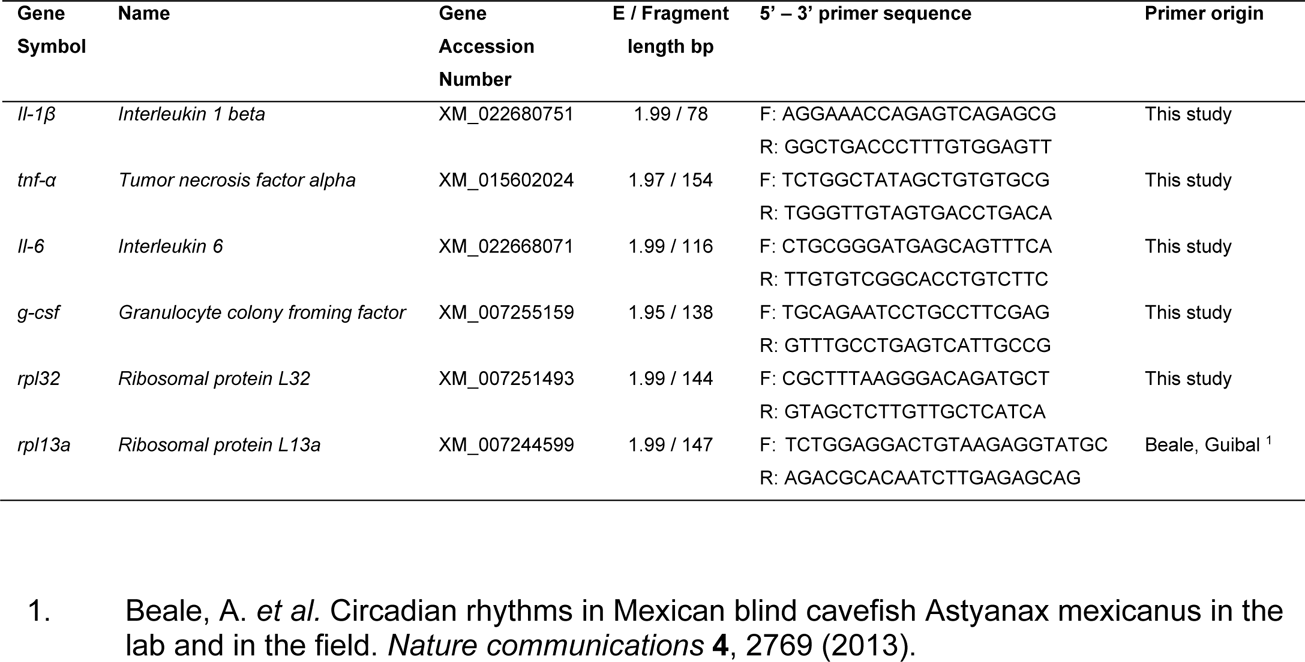
Primer-sequences with respective efficiencies (E) for RT-qPCR.

**Table S2:** Overview of features extracted for morphological identification of A. mexicanus head kidney cells.

| Feature ID | Feature_ImageMask_Channel | Feature description |
| --- | --- | --- |
| 1 | Area_AdaptiveErode_BF | Cell size |
| 2 | Area_Intensity_SSC | Areas of SSC signal above background |
| 3 | Area_Morphology_DNA | Area of DNA signal (nuclear staining) |
| 4 | Aspect.Ratio_AdaptiveErode_BF | Aspect ratio of total cell area |
| 5 | Bright.Detail.Intensity.R3_AdaptiveErode_BF_BF | Intensity of brightest staining areas |
| 6 | Bright.Detail.Intensity.R3_AdaptiveErode_BF_Draq5 | Intensity of brightest staining areas |
| 7 | Bright.Detail.Intensity.R3_AdaptiveErode_BF_SSC | Intensity of brightest signal areas |
| 8 | Circularity_AdaptiveErode_BF | Circularity of whole cell shape |
| 9 | Circularity_Morphology_DNA | Circularity of nucleus |
| 10 | Contrast_AdaptiveErode_BF_BF | Detects large changes in pixel values - can be measure of granularity of signal |
| 11 | Contrast_AdaptiveErode_BF_SSC | Detects large changes in pixel values - can be measure of granularity of signal |
| 12 | Diameter_AdaptiveErode_BF | Diameter of whole cell shape |
| 13 | Diameter_Morphology_DNA | Diameter of nucleus |
| 14 | H.Energy.Mean_AdaptiveErode_BF_BF_5 | Measure of intensity concentration - texture feature |
| 15 | H.Energy.Mean_Morphology_DNA_Draq5_5 | Measure of intensity concentration - texture feature |
| 16 | H.Entropy.Mean_AdaptiveErode_BF_BF_5 | Measure of intensity concentration and randomness of signal - texture feature |
| 17 | H.Entropy.Mean_Morphology_DNA_Draq5_5 | Measure of intensity concentration and randomness of signal - texture feature |
| 18 | Intensity_AdaptiveErode_BF_Draq5 | Integrated intensity of signal within whole cell mask |
| 19 | Intensity_AdaptiveErode_BF_SSC | Integrated intensity of signal within whole cell mask |
| 20 | Lobe.Count_Morphology_DNA_Draq5 | Number of lobes of nucleus |
| 21 | Max.Pixel_Intensity_Ch6_SSC | maximum pixel intensity of stated channel within a whole cell mask |
| 22 | Max.Pixel_Morphology_DNA_Draq5 | maximum pixel intensity of stated channel within a whole cell mask |
| 23 | Mean.Pixel_Morphology_DNA_Draq5 | mean pixel intensity of stated channel within a whole cell mask |
| 24 | Shape.Ratio_AdaptiveErode_BF | minimum thickness divided by length - measure of cell shape characteristic |
| 25 | Std.Dev_AdaptiveErode_BF_BF | standard deviation of BF signal - measure of granularity and variance in BF |

