## Supplementary material for "Adaptation to low parasite abundance affects immune investment strategy and immunopathological responses of cavefish": Data File 2_Cell Gallery Cluster

### Slide 1
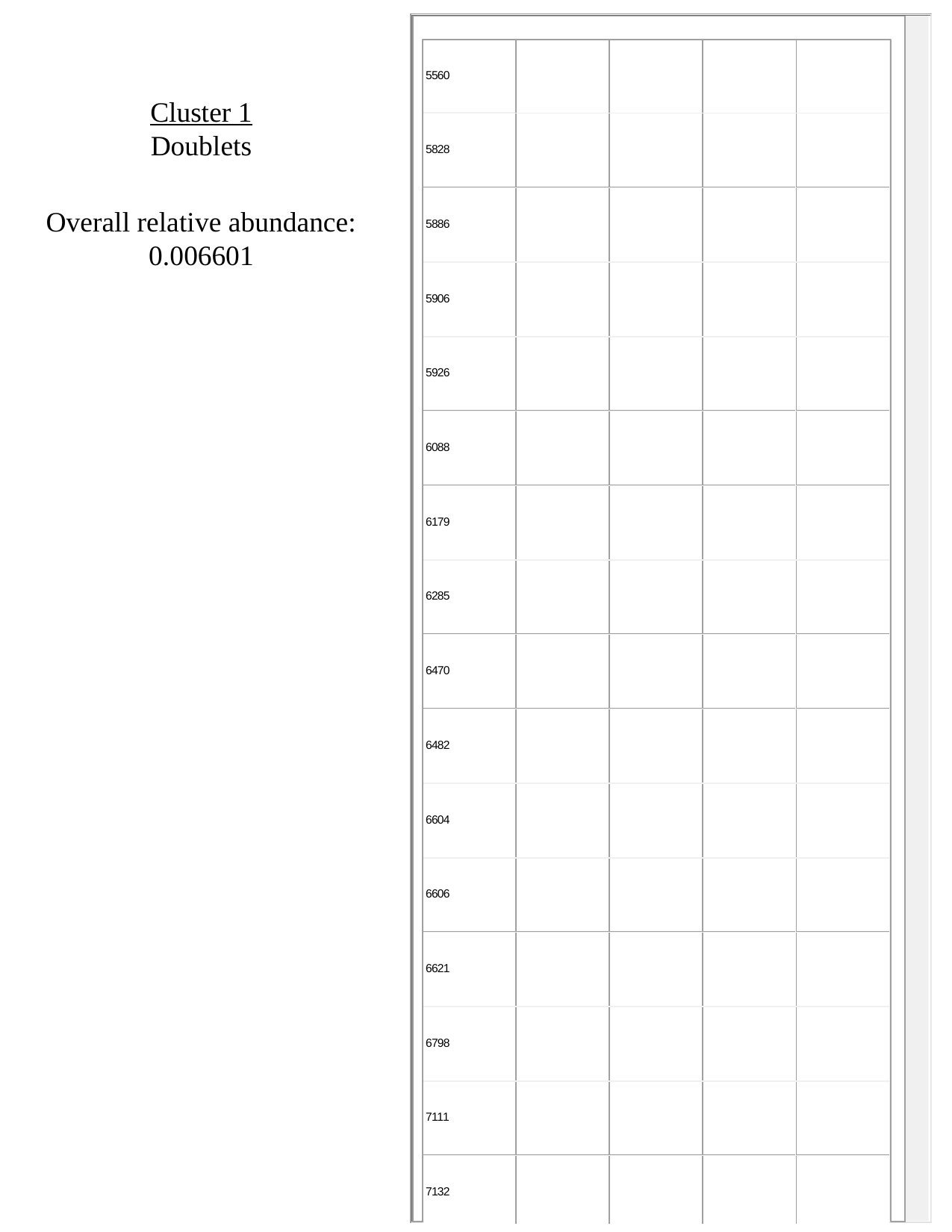

Cluster 1
Doublets
Overall relative abundance:
0.006601

### Slide 2
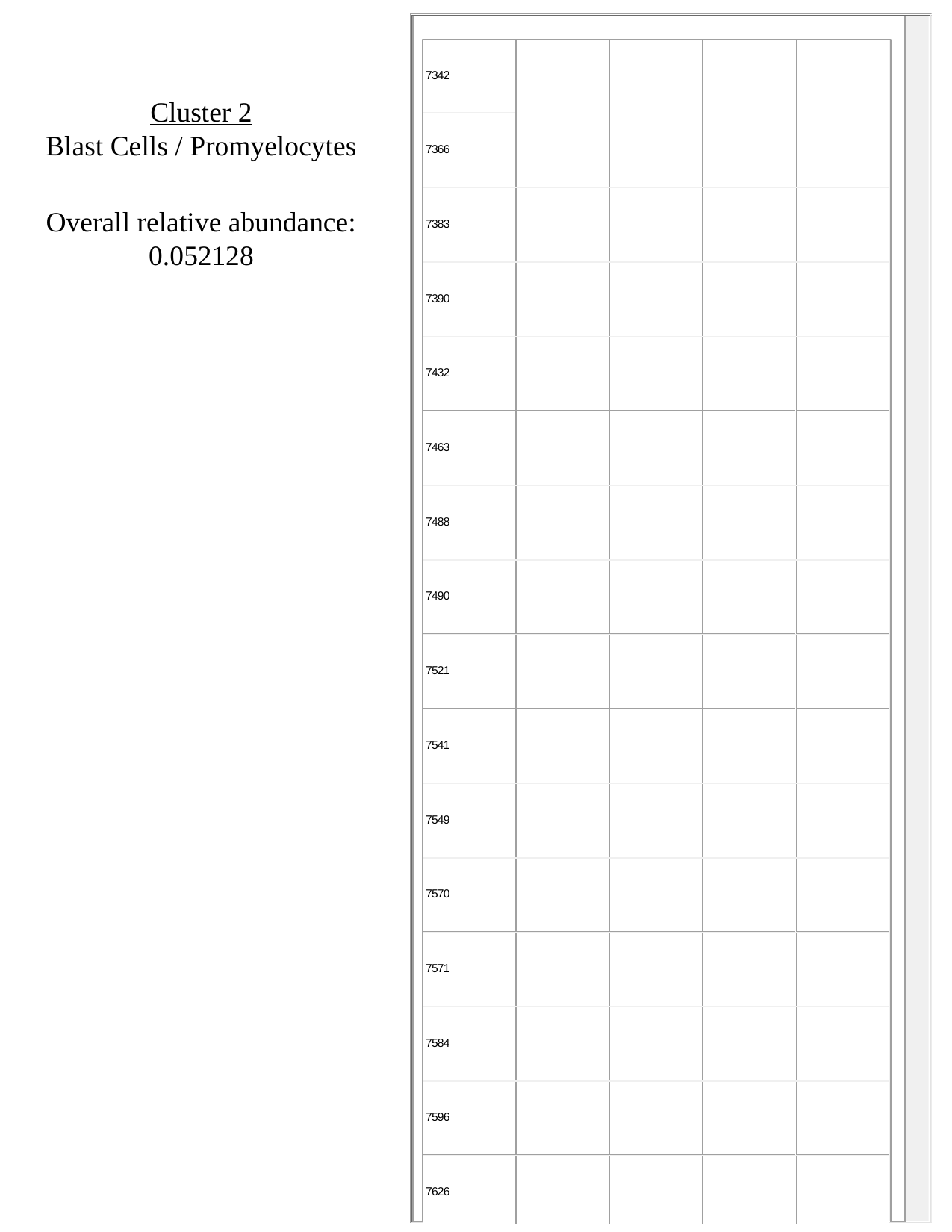

Cluster 2
Blast Cells / Promyelocytes
Overall relative abundance:
0.052128

### Slide 3
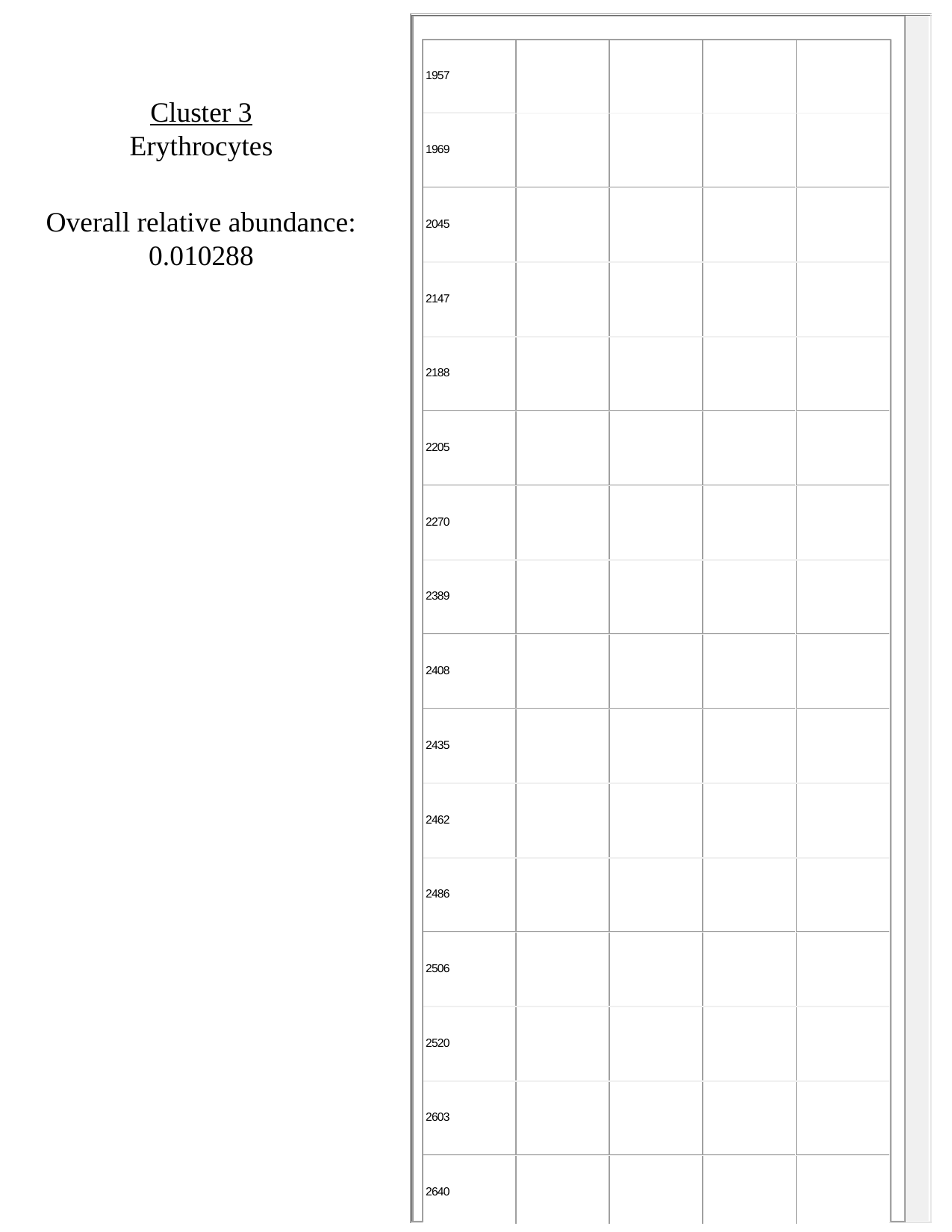

Cluster 3
Erythrocytes
Overall relative abundance:
0.010288

### Slide 4
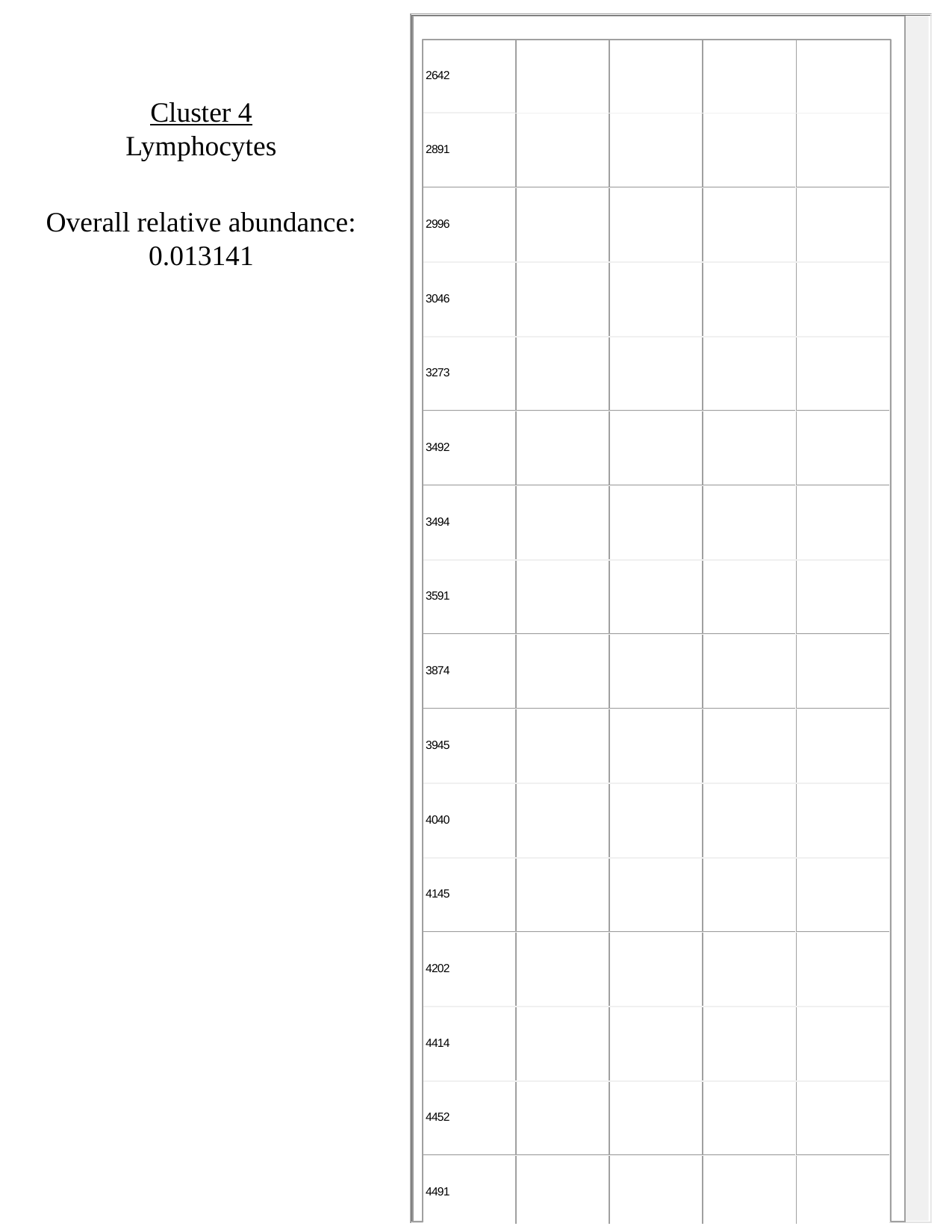

Cluster 4
Lymphocytes
Overall relative abundance:
0.013141

### Slide 5
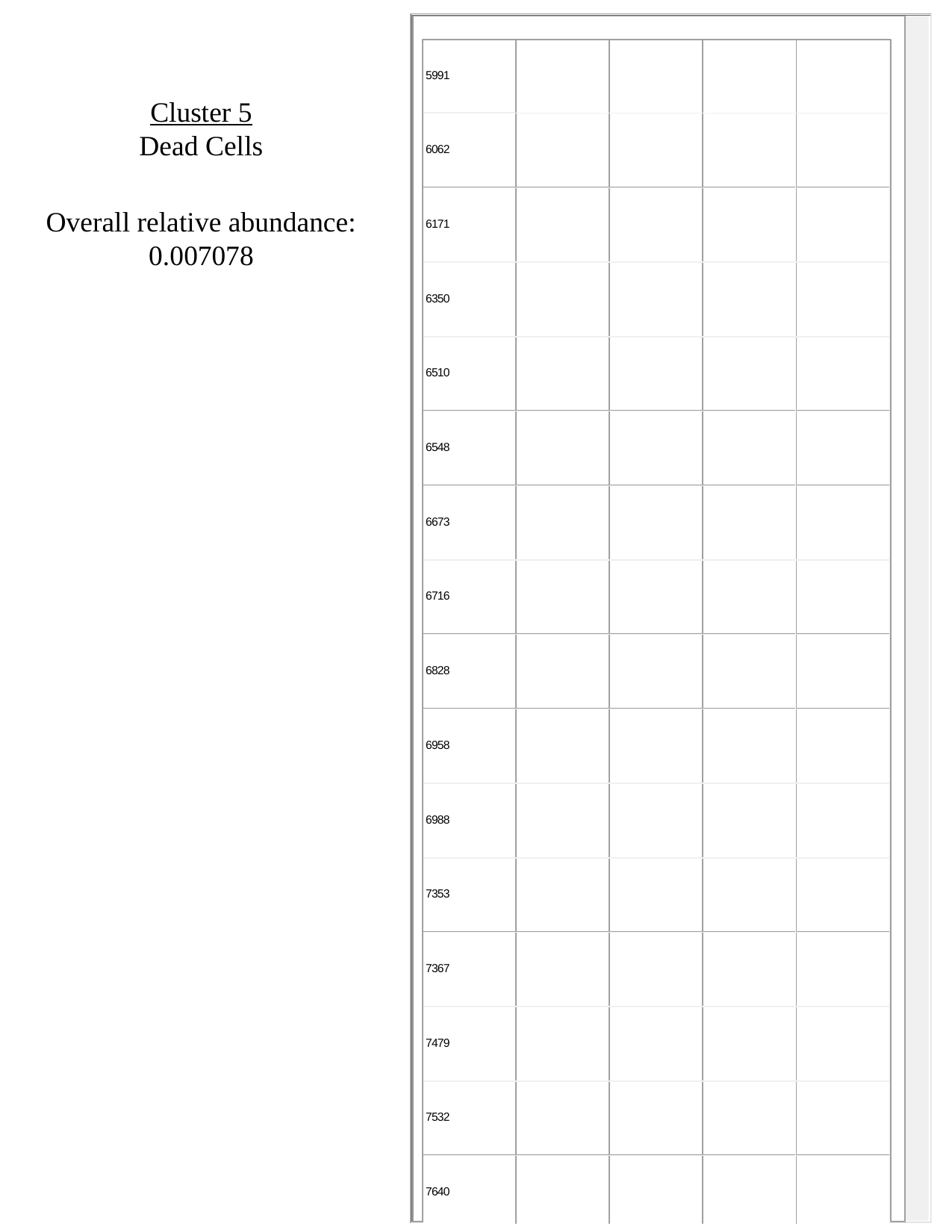

Cluster 5
Dead Cells
Overall relative abundance:
0.007078

### Slide 6
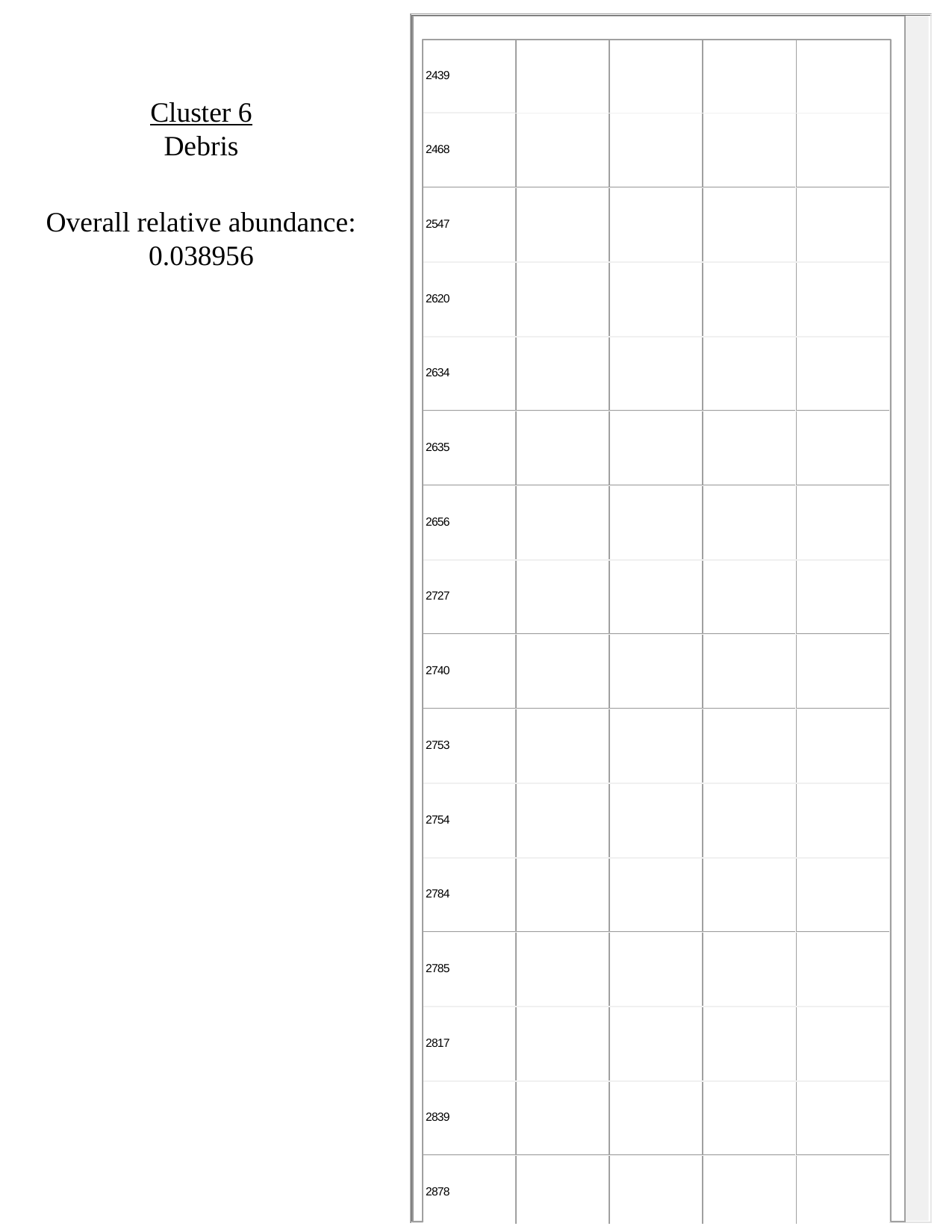

Cluster 6
Debris
Overall relative abundance:
0.038956

### Slide 7
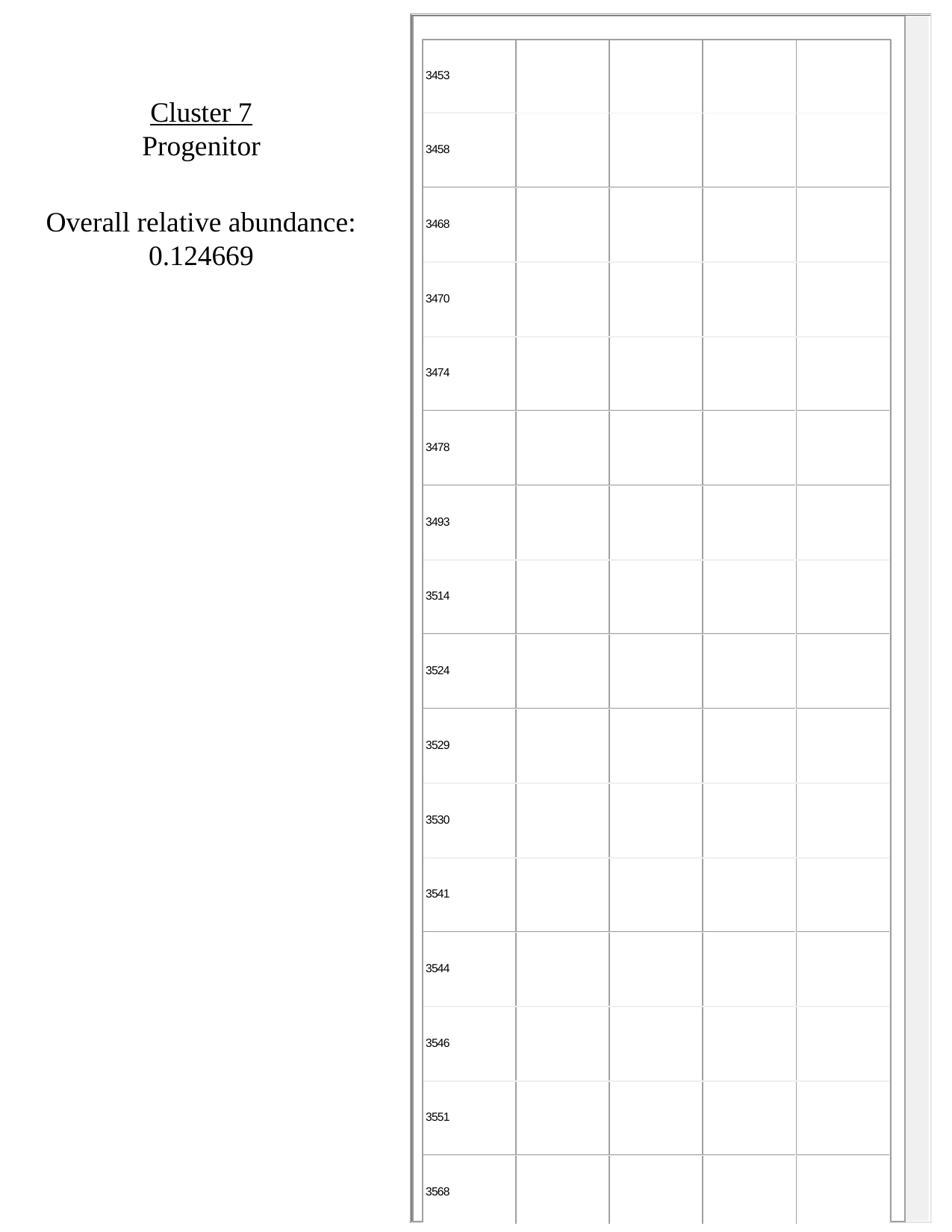

Cluster 7
Progenitor
Overall relative abundance:
0.124669

### Slide 8
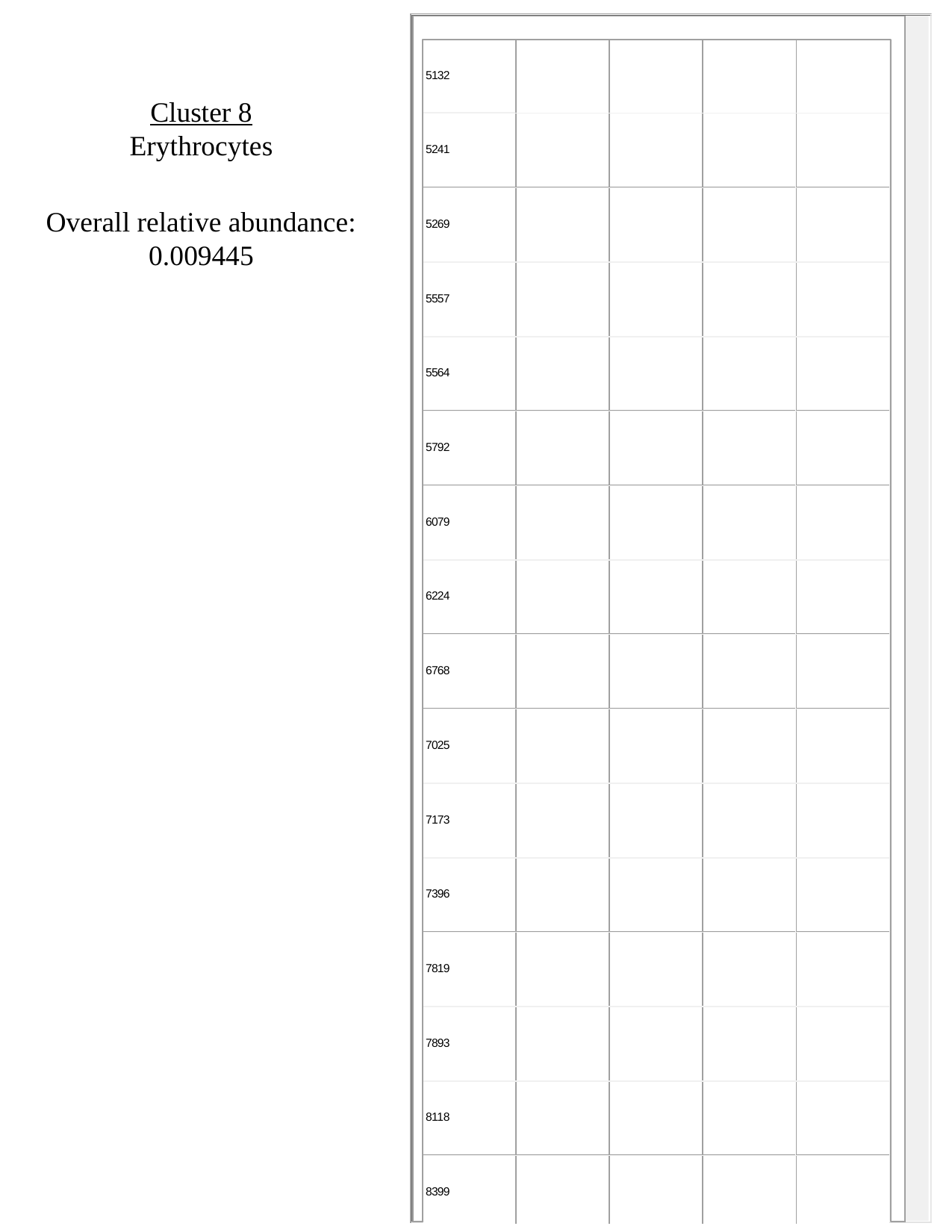

Cluster 8
Erythrocytes
Overall relative abundance:
0.009445

### Slide 9
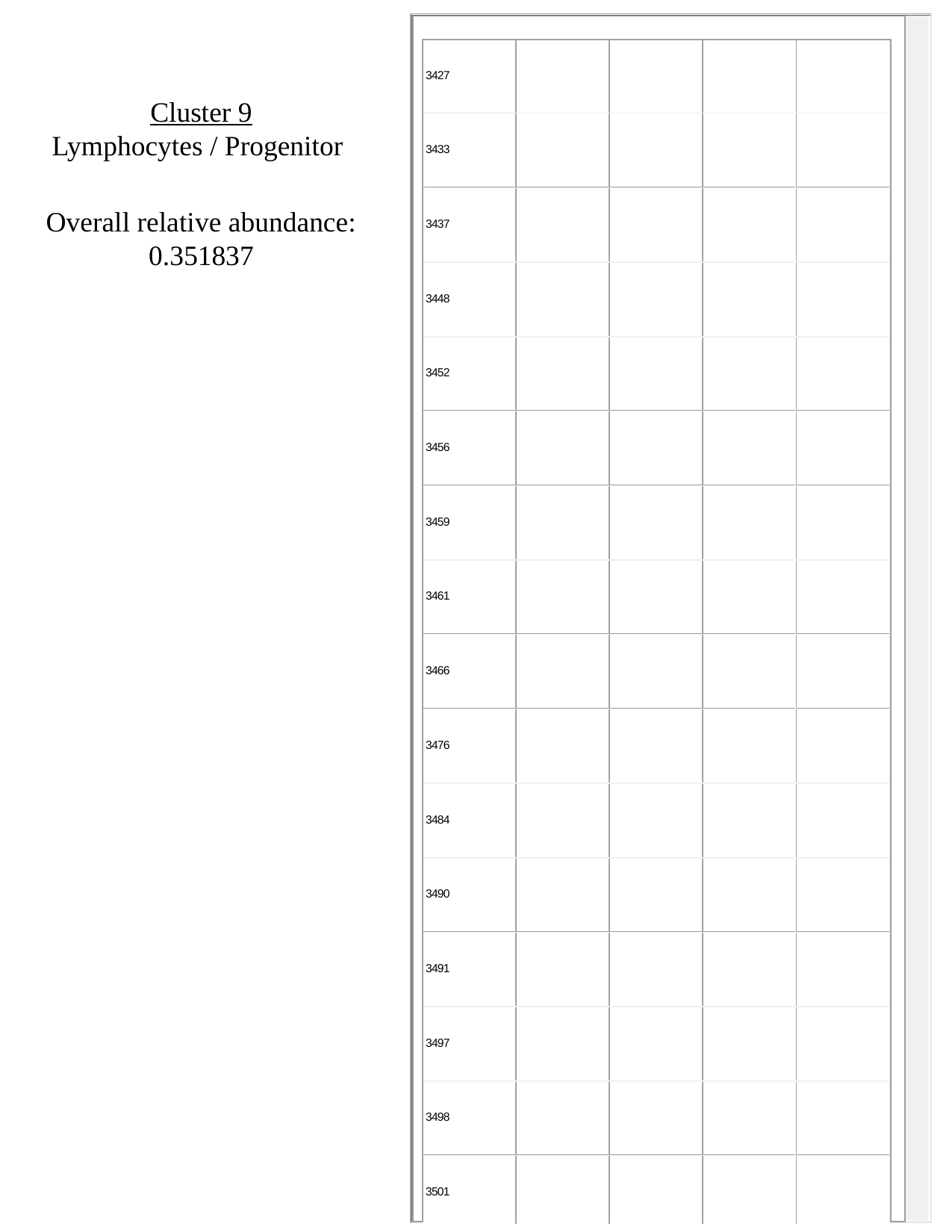

Cluster 9
Lymphocytes / Progenitor
Overall relative abundance:
0.351837

### Slide 10
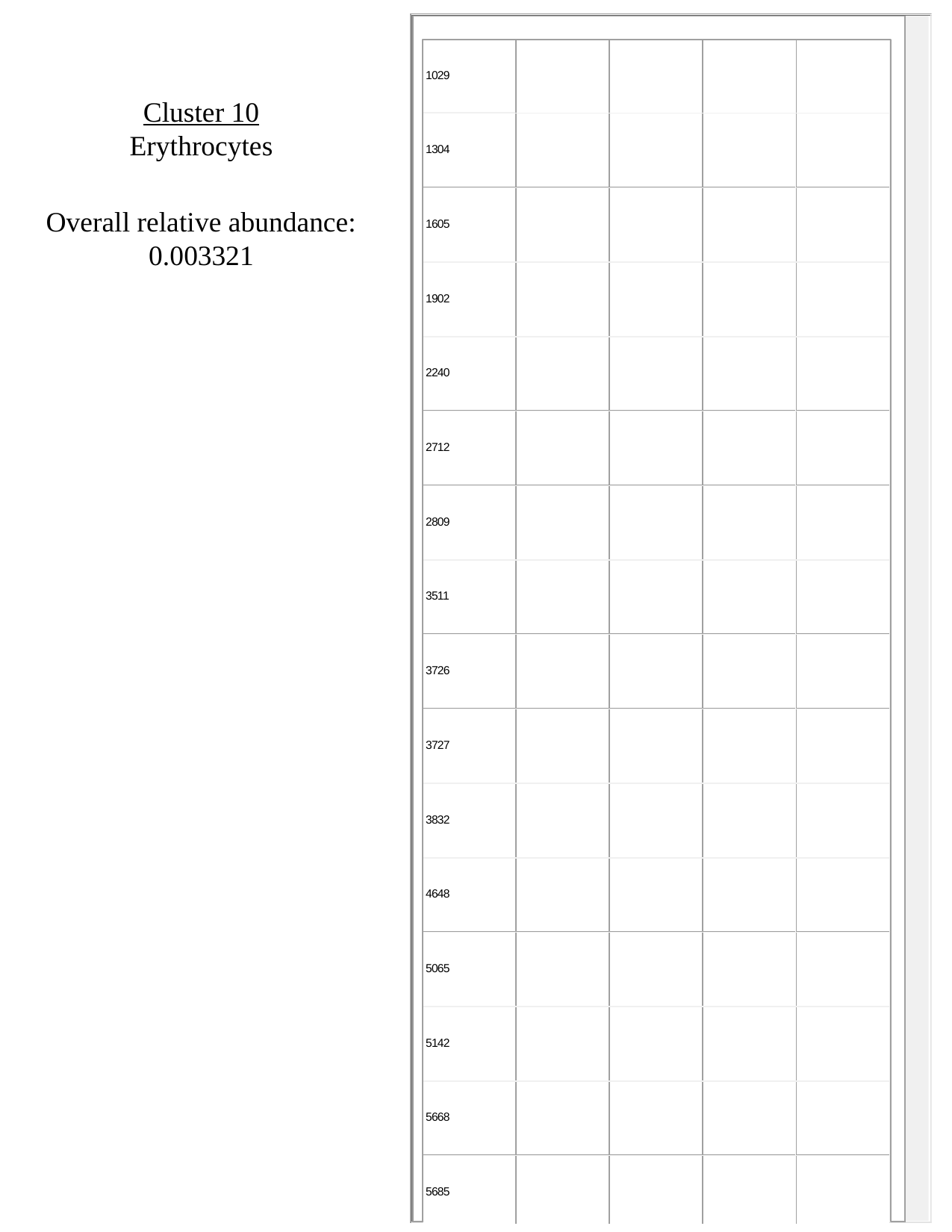

Cluster 10
Erythrocytes
Overall relative abundance:
0.003321

### Slide 11
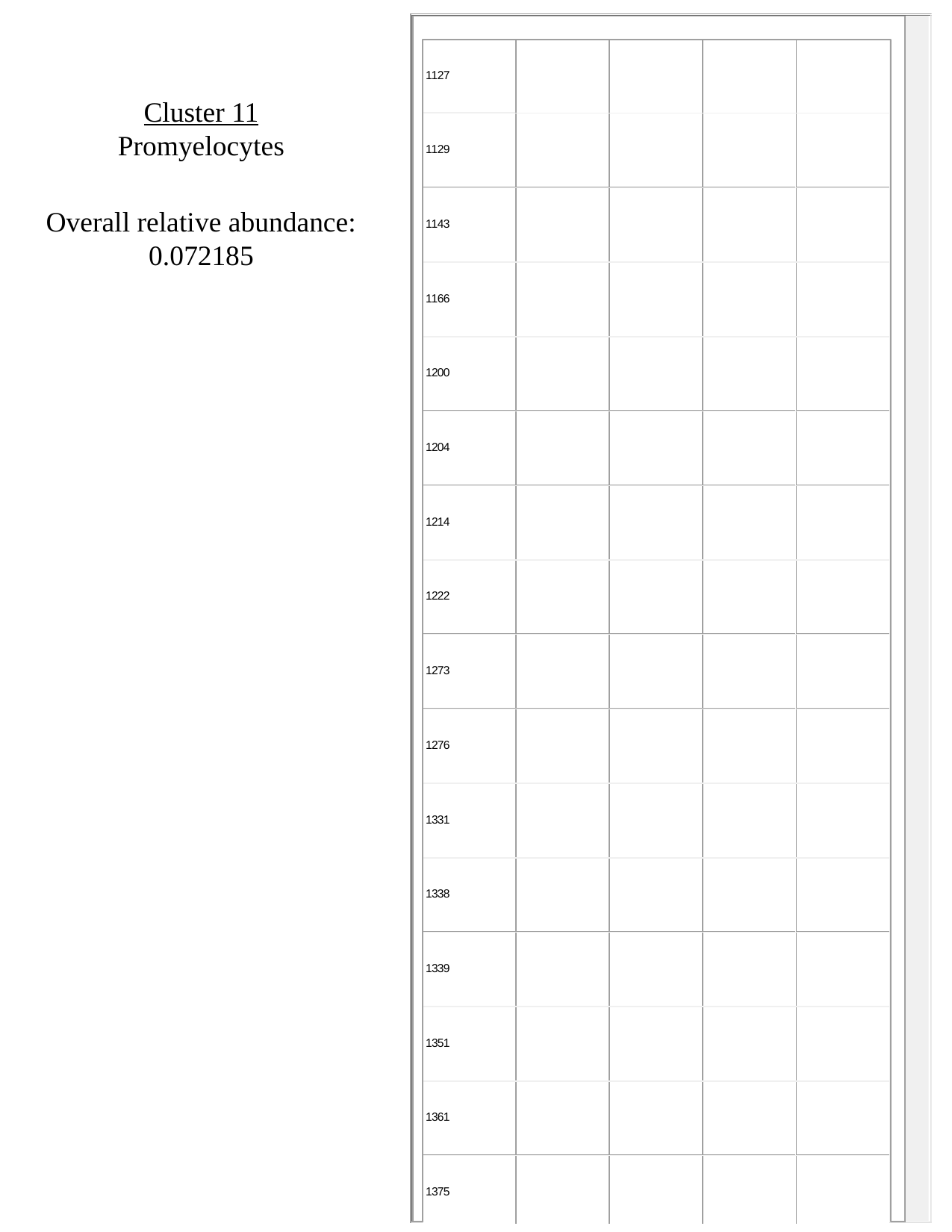

Cluster 11
Promyelocytes
Overall relative abundance:
0.072185

### Slide 12
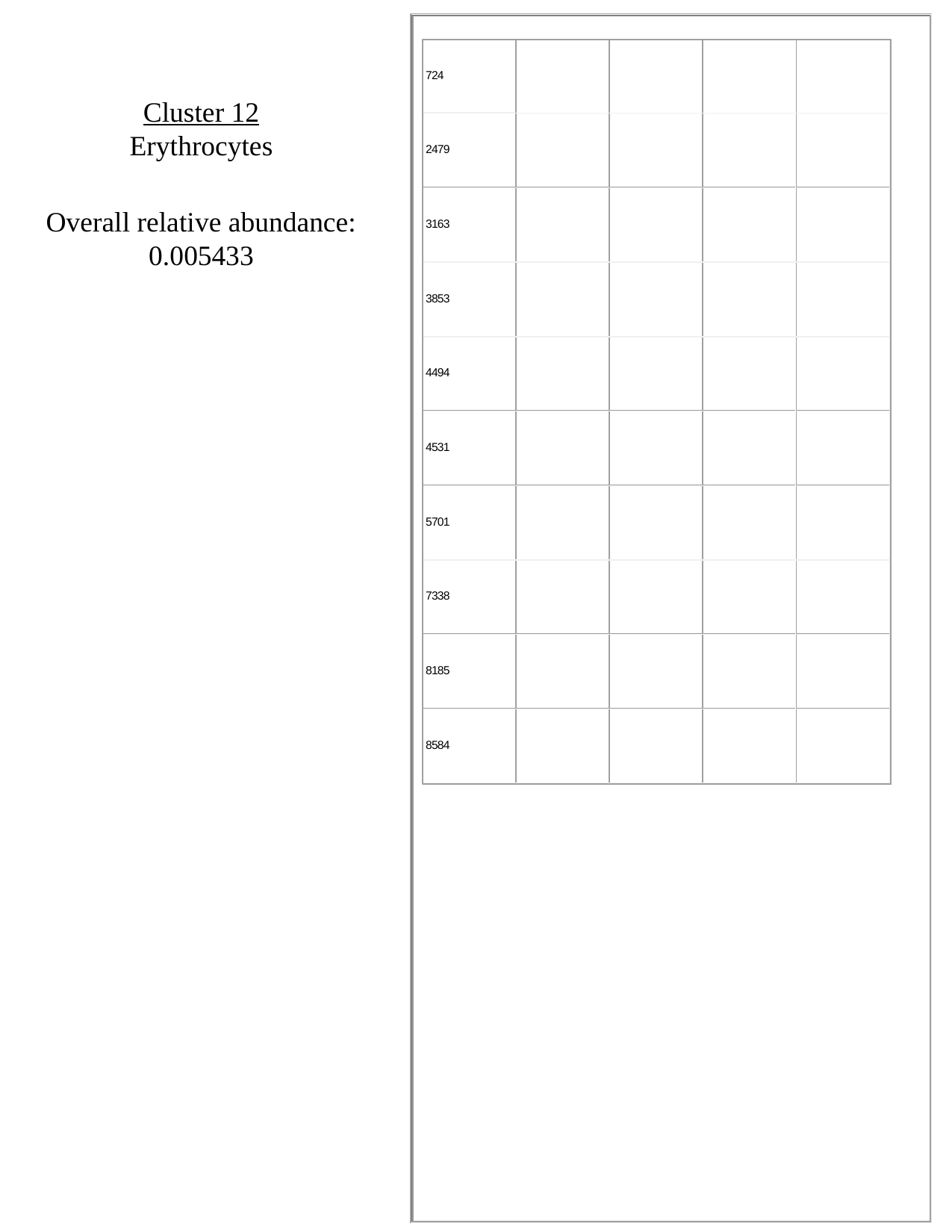

Cluster 12
Erythrocytes
Overall relative abundance:
0.005433

### Slide 13
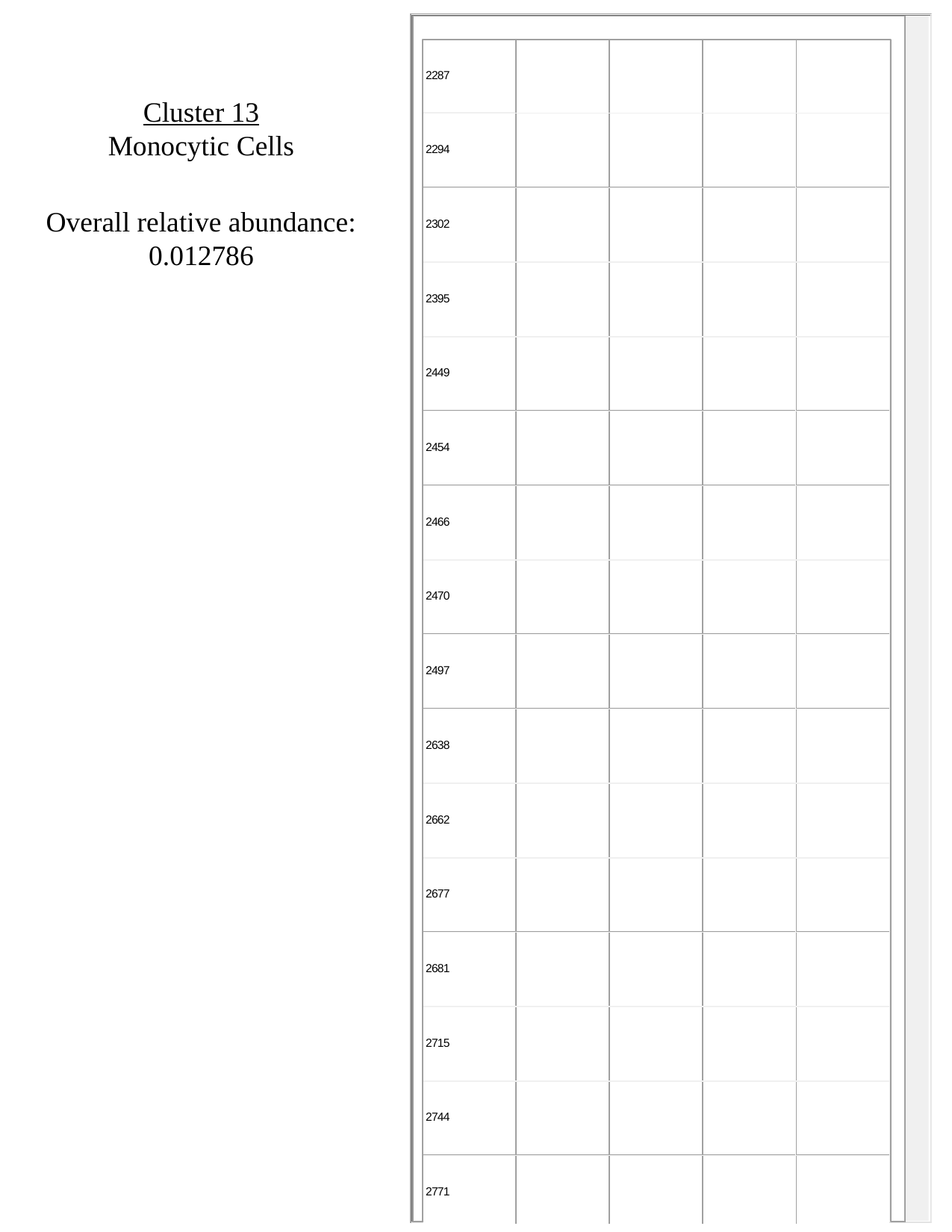

Cluster 13
Monocytic Cells
Overall relative abundance:
0.012786

### Slide 14
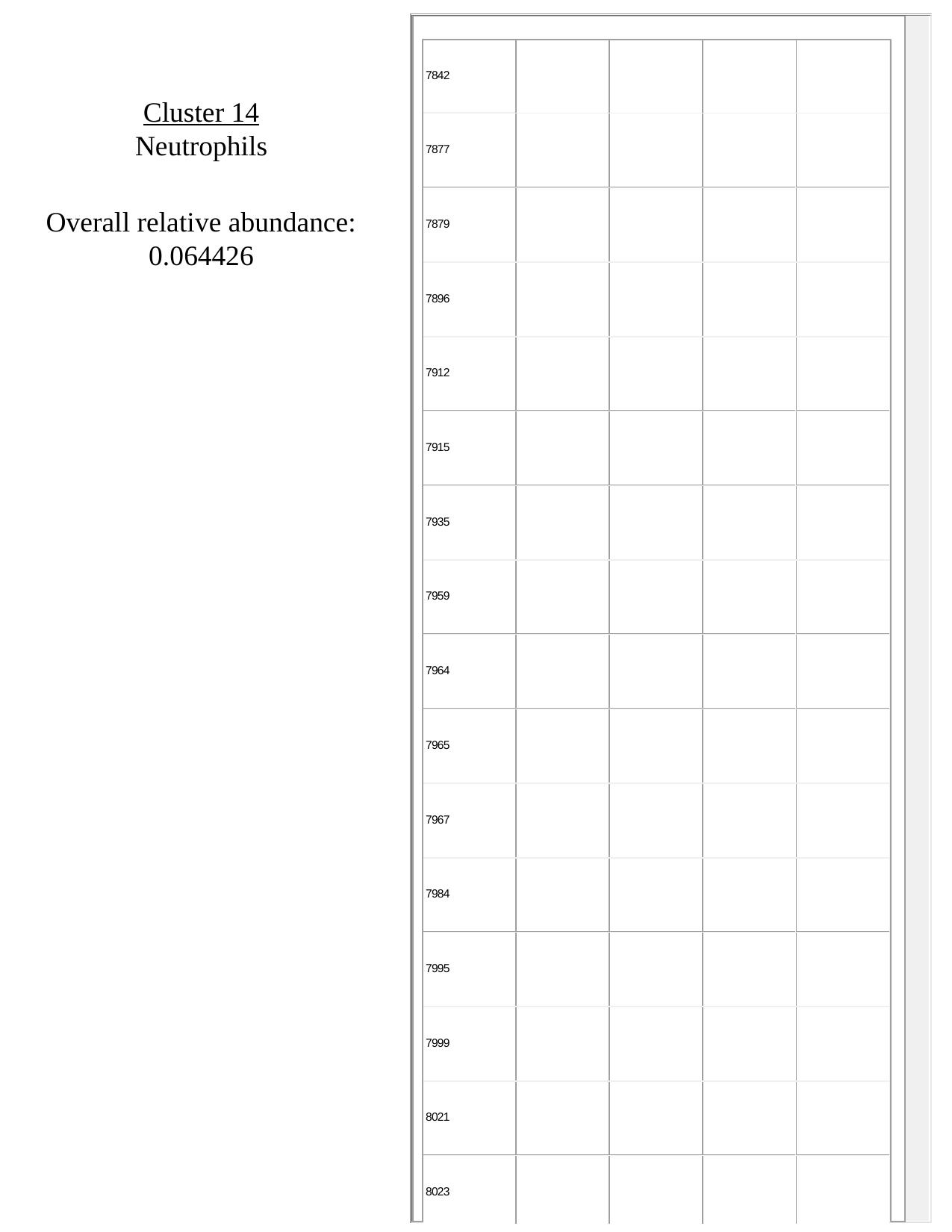

Cluster 14
Neutrophils
Overall relative abundance:
0.064426

### Slide 15
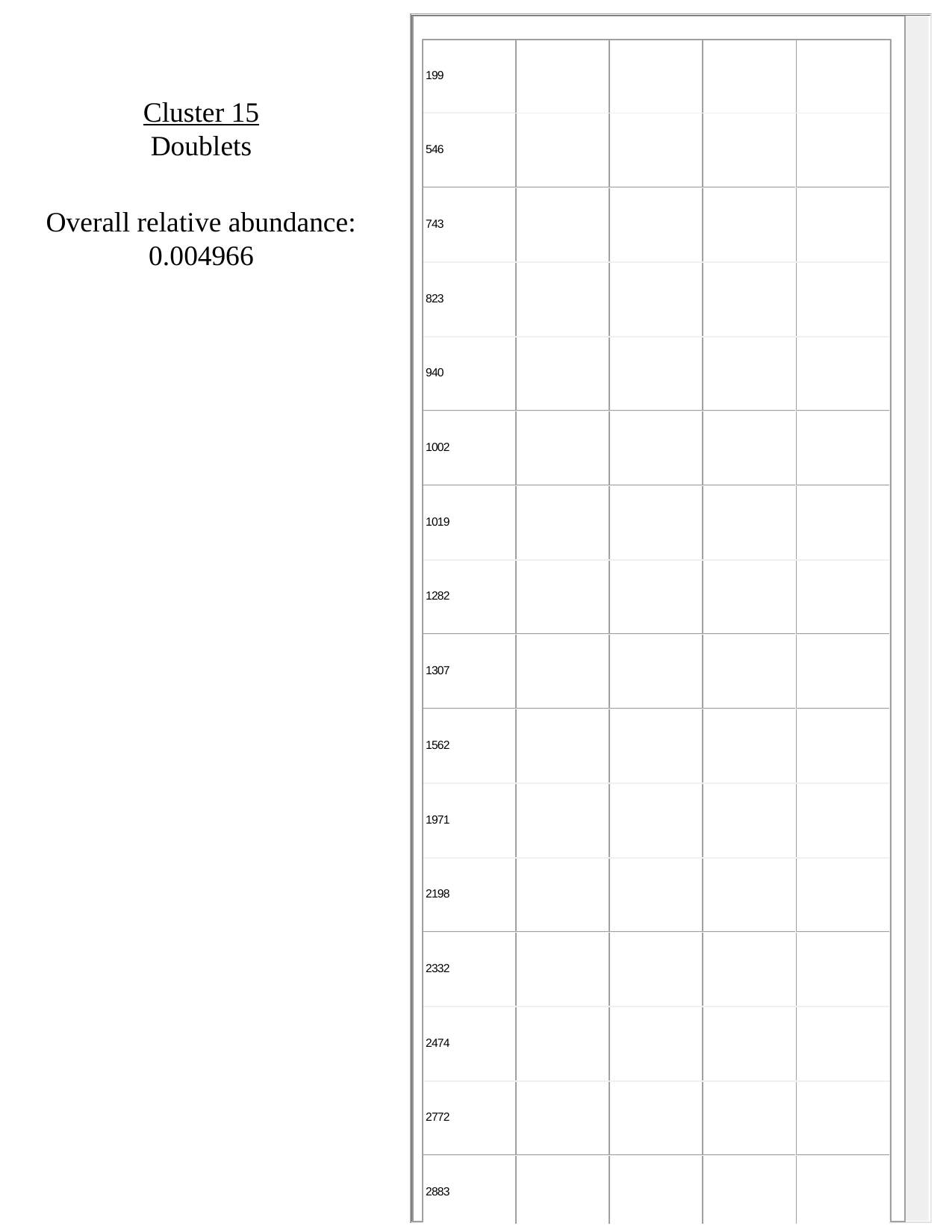

Cluster 15
Doublets
Overall relative abundance:
0.004966

### Slide 16
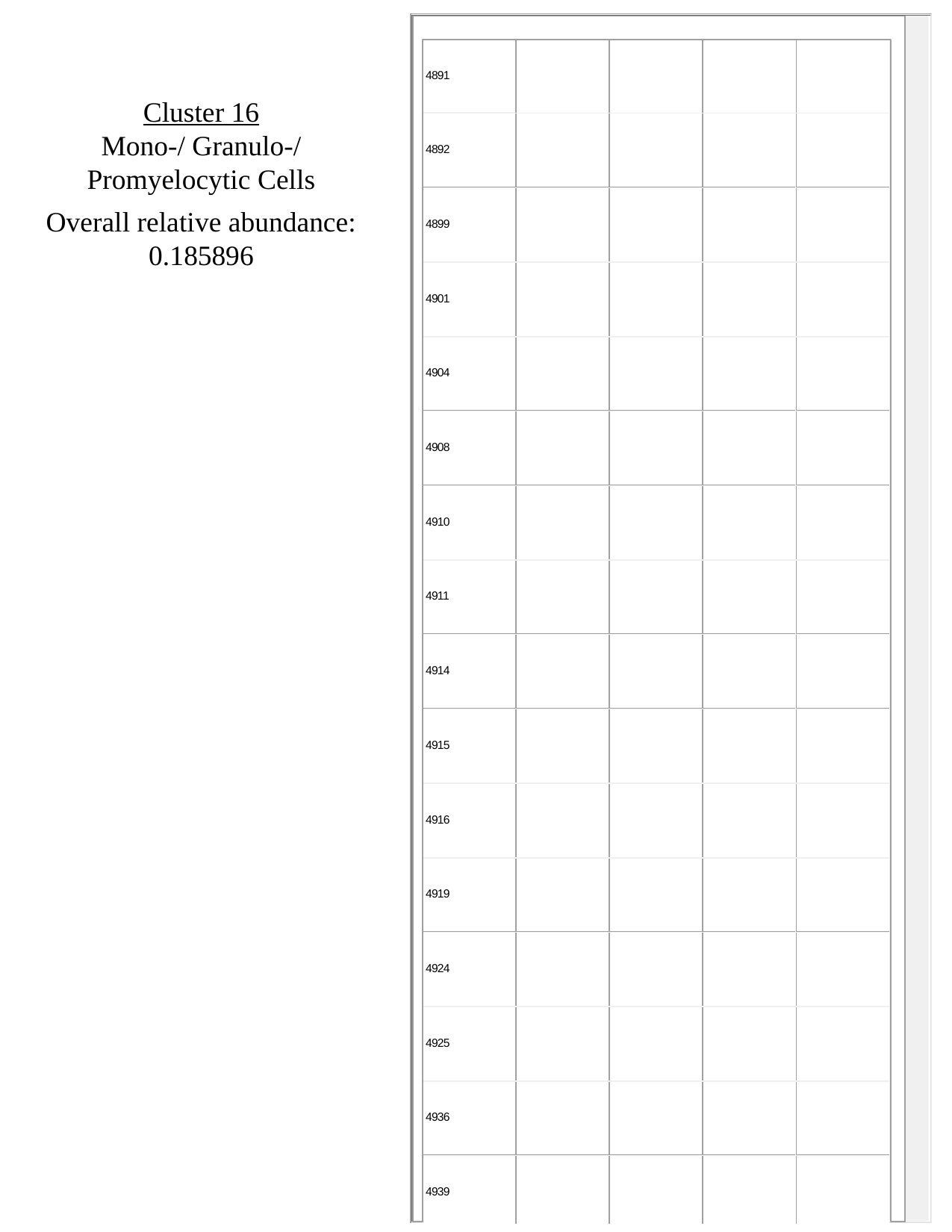

Cluster 16
Mono-/ Granulo-/ Promyelocytic Cells
Overall relative abundance:
0.185896

### Slide 17
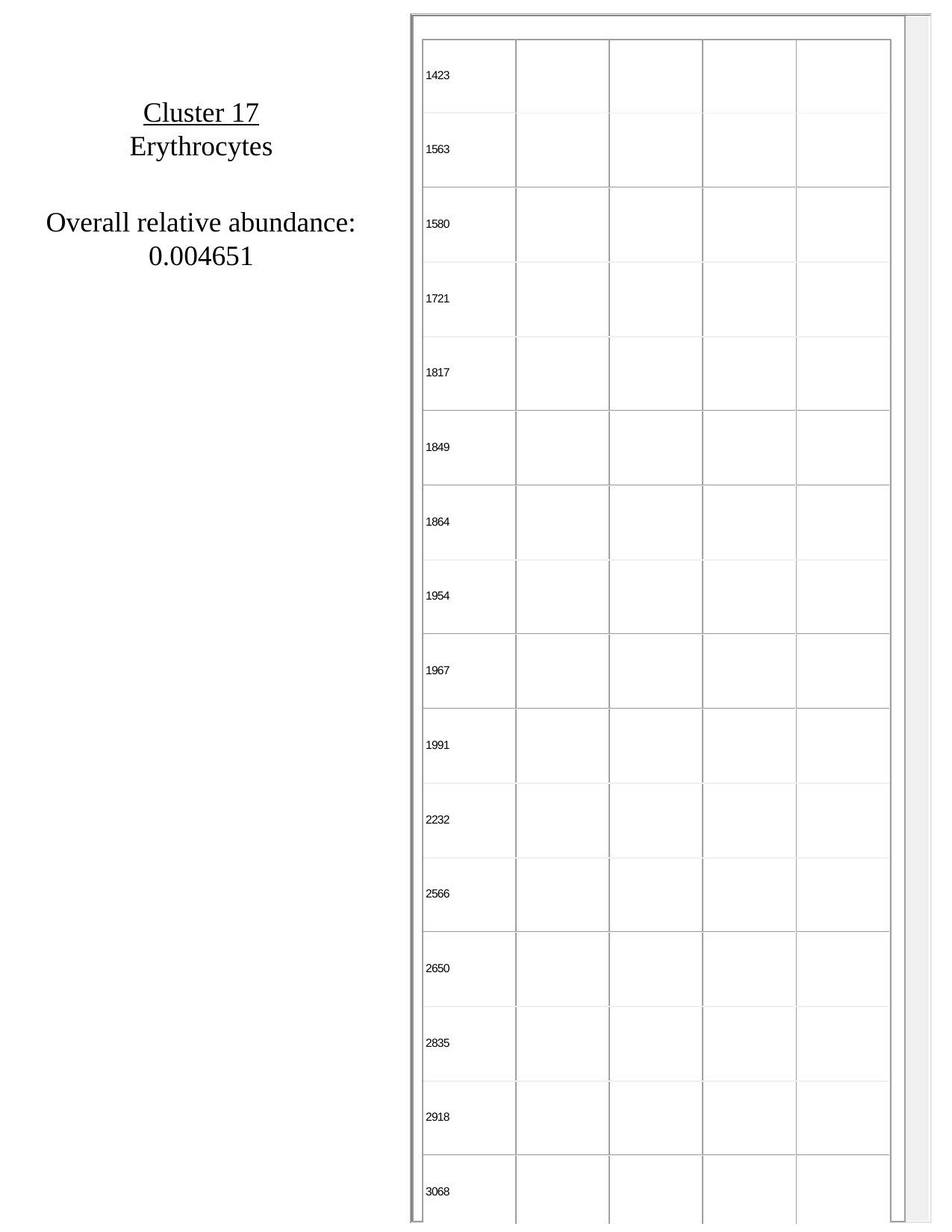

Cluster 17
Erythrocytes
Overall relative abundance:
0.004651

### Slide 18
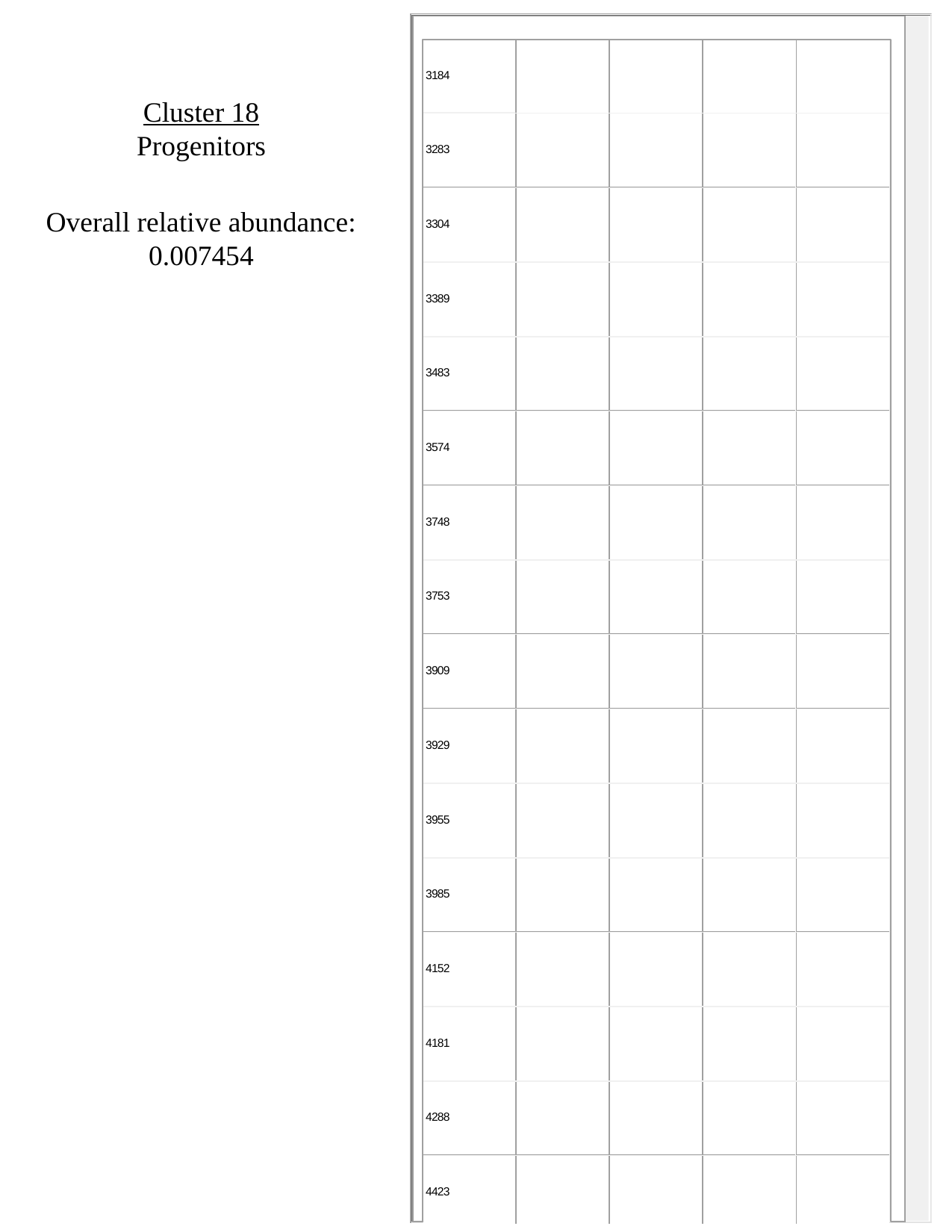

Cluster 18
Progenitors
Overall relative abundance:
0.007454

### Slide 19
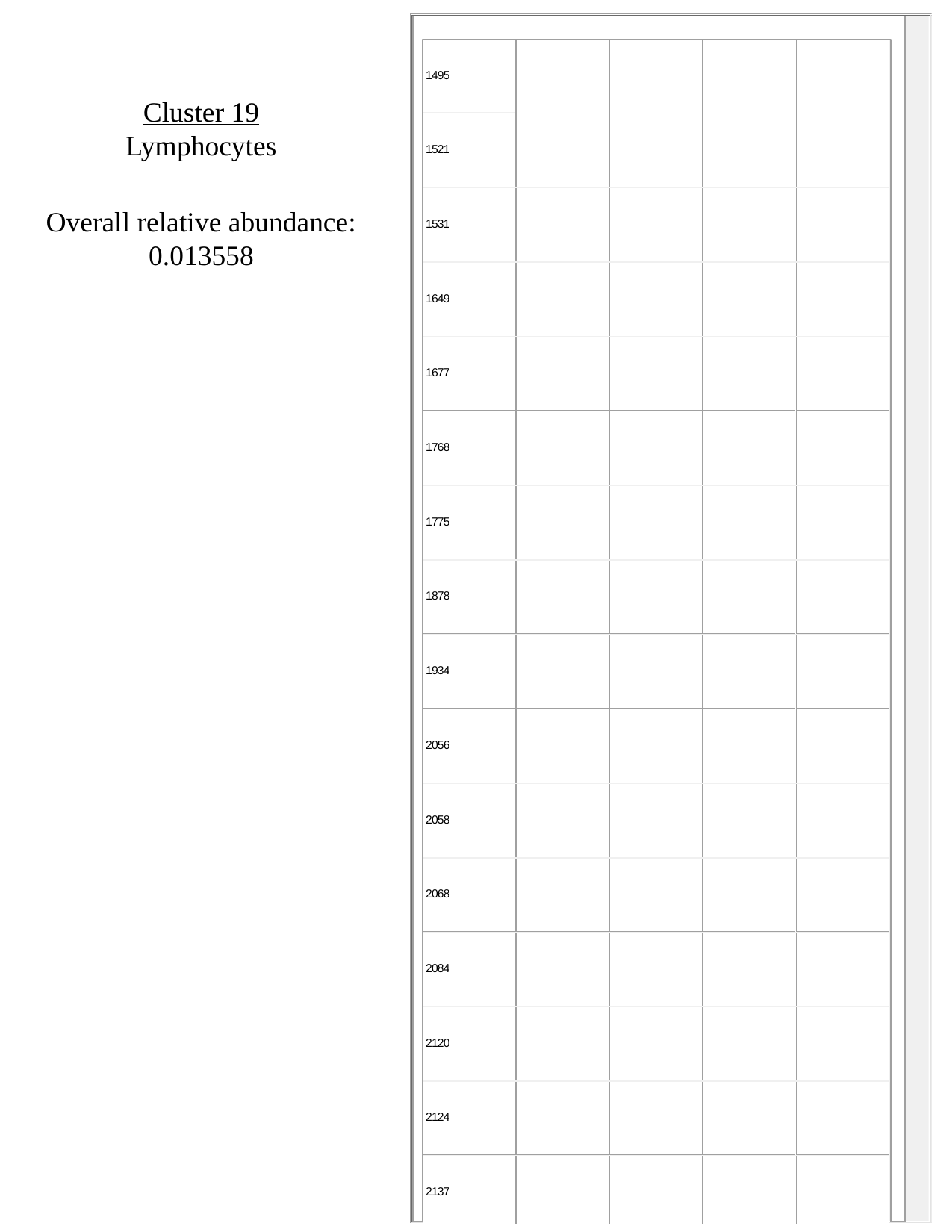

Cluster 19
Lymphocytes
Overall relative abundance:
0.013558

### Slide 20
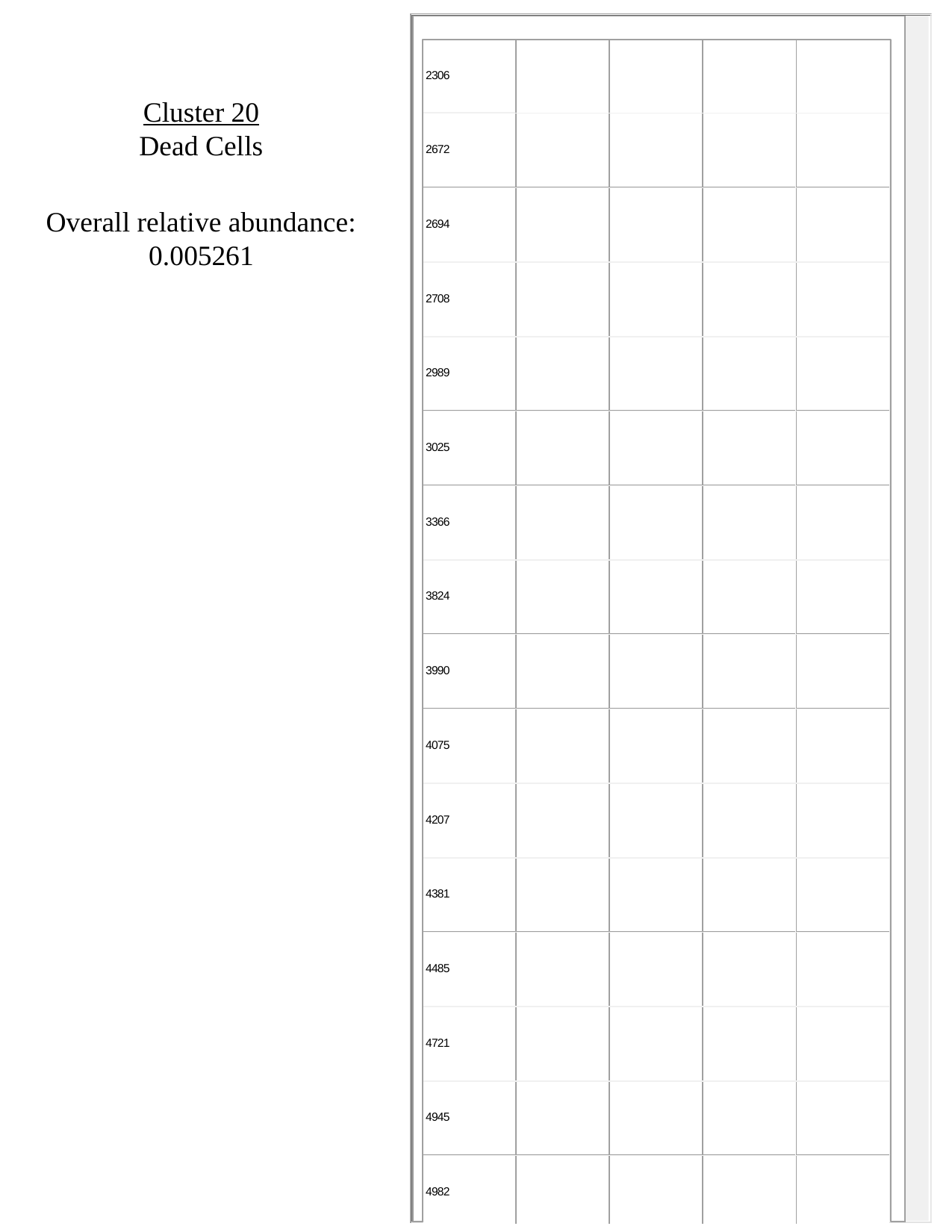

Cluster 20
Dead Cells
Overall relative abundance:
0.005261

### Slide 21
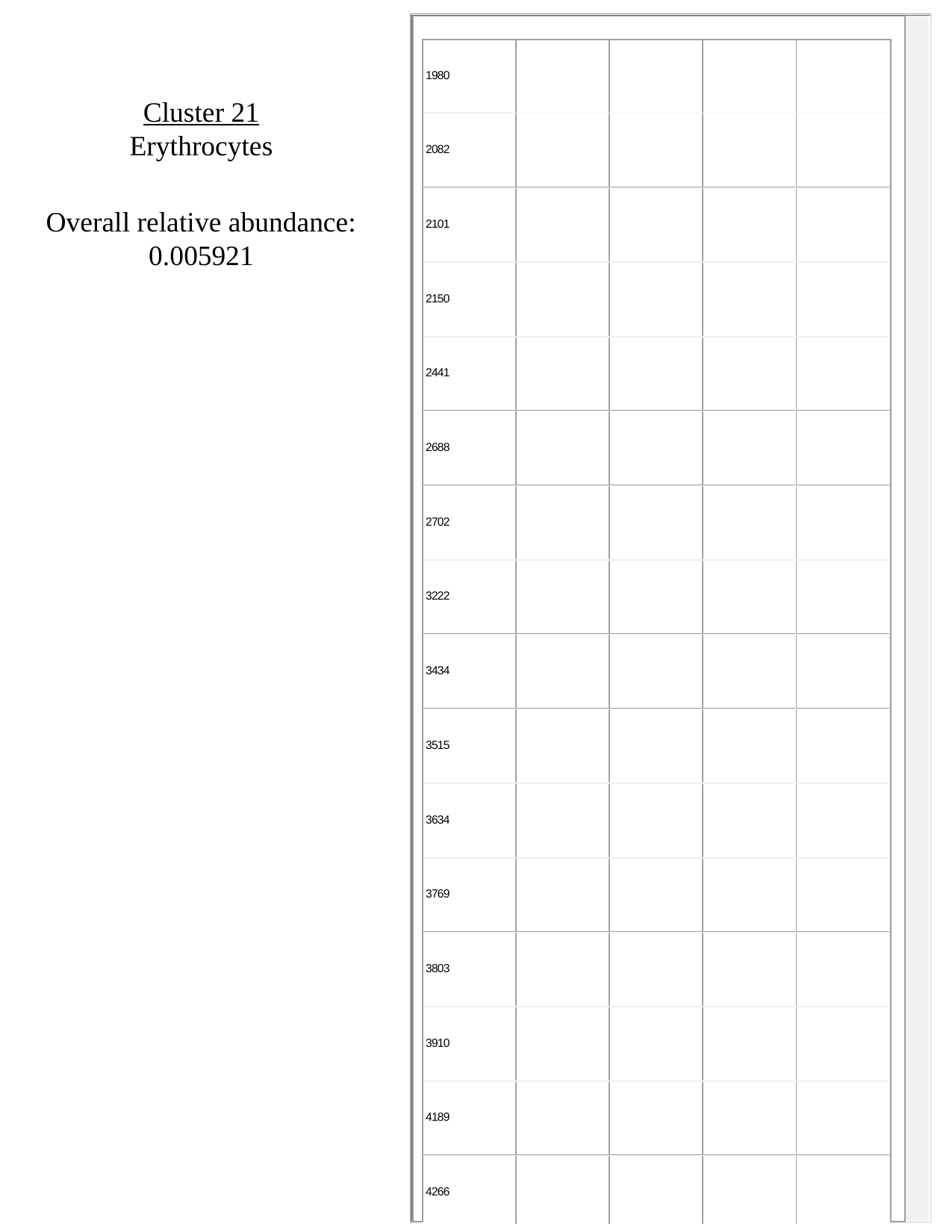

Cluster 21
Erythrocytes
Overall relative abundance:
0.005921
