## Supplementary figures and images for "Adaptation to low parasite abundance affects immune investment strategy and immunopathological responses of cavefish"

### Data File 5_SurfacePBSMarkerHeatmap

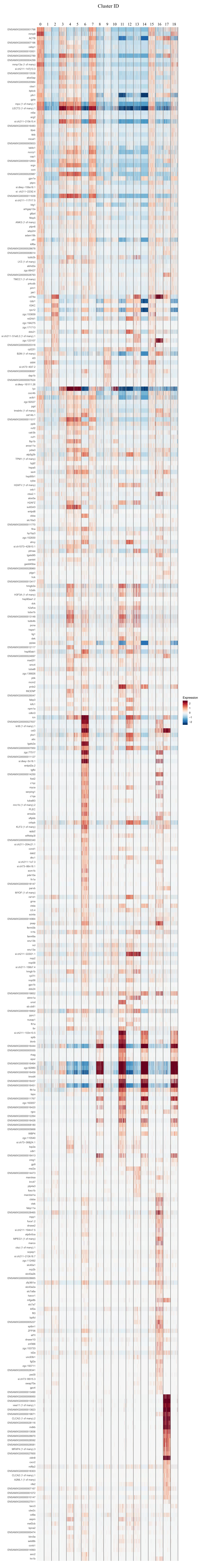

### Data File 6_SurfaceLPSMarkerHeatmap

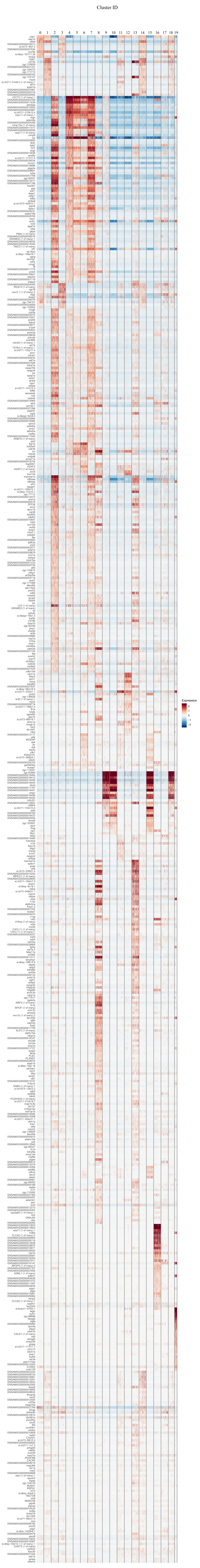

### Data File 7_CavefishPBSMarkerHeatmap

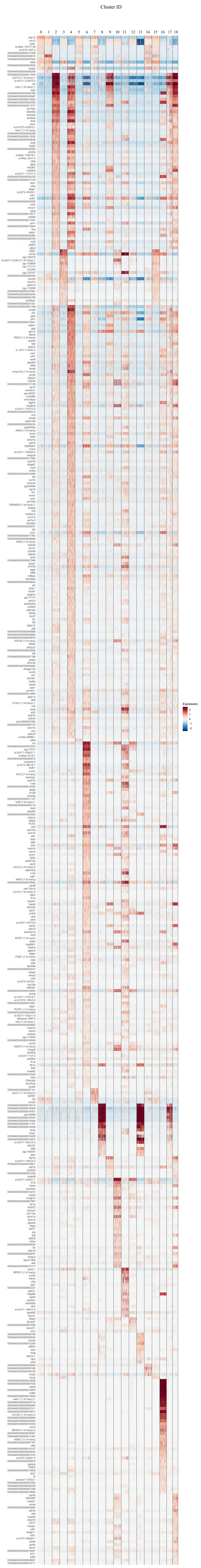

### Data File 8_CavefishLPSMarkerHeatmap

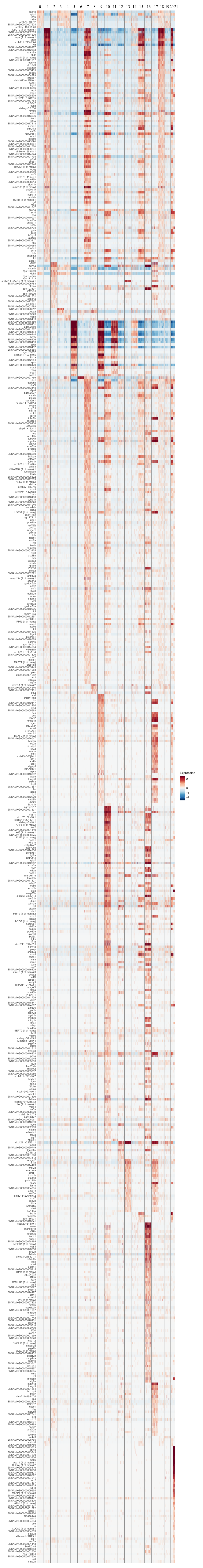
